# Determining the scale at which variation in WUE traits changes population yields

**DOI:** 10.1101/682674

**Authors:** Erica McGale, Henrique Valim, Deepika Mittal, Jesús Morales Jimenez, Rayko Halitschke, Meredith C. Schuman, Ian T. Baldwin

## Abstract

Functional variation is known to influence population yield, but the scale at which this happens is still unknown. Relevant signals might only reach immediate neighbors of a phenotypically diverse plant (neighbor-scale) or conversely may distribute across the population (population-scale). We use *Nicotiana attenuata* silenced in mitogen-activated protein kinase 4 (irMPK4), plants with low water-use efficiency (WUE), to study the scale at which water-use traits alter intraspecific population yields. In the field and glasshouse, populations with low percentages of irMPK4 plants planted among isogenic control plants produced maximum overall growth and yield. Through paired-plant and local-plant-configuration analyses, we determined that this occurred at the population scale. However, we find that this effect was not due to irMPK4’s WUE phenotype. With micro-grafting, we additionally show that *MPK4*-deficiency may mediate the response at the population-scale: shoot-expressed *MPK4* is required for *N. attenuata* to change yield in response to a neighbor.

## Introduction

Scientists express biological complexity as hierarchical levels in order to provide structure for the study of biological phenomena. By definition, higher levels in biological hierarchies set boundaries for the function of traits at lower levels in the hierarchy, but do not define the function of these traits (Allen and Starr, 1982). In plant ecology, an individual plant represents a lower organizational level of a population while a population represents a lower level of a community. Within a population, variation in trait expression can alter the function of individuals, which in turn can create significant changes at the population level. Total population growth and yield (defined here as biomass and fitness correlates, i.e. flowers and seed capsules) is influenced by the diversity of traits expressed by the individuals in a population, but the mechanisms responsible for these population yield changes are not known, and many feed-back loops which exist at different scales in the hierarchy have been proposed. Within a population, individual plants could integrate information either about total population performance (i.e. at the population scale) or immediate neighbor performance (i.e. at the neighbor scale) and respond to this information by changing their individual growth and yield. At the population-scale, responding individuals (RIs) might interpret the cumulative state of signals (defined here as information in transit among individuals with the ability to communicate; Allen and Starr, 1982) in a population. RIs would then respond homogenously across a population, causing changes in population yield in direct proportion to RI abundance in the population (Crawford and Rudgers, 2012; Smith and Knapp, 2003). However, RIs might also communicate with immediate neighbors through signals at the neighbor scale, creating heterogeneous changes in yield across the population wherever RIs and specific neighbor types interact. Neighbor-scale responses do not require the context of the population to contribute to changes in local yields, but the sums of these local yields can change population yield (Crawford and Rudgers, 2012; Hughes et al., 2008).

Whether interpreted at the neighbor or population scale, signals which cause yield responses can originate from water-use trait variation within populations: for example, variation in water-use efficiency (WUE) of individuals is known to alter population growth and yield (Caldeira et al., 2001; Comas et al., 2013; Forrester, 2015; Kimball et al., 2014; Marguerit et al., 2014; Wang et al., 2016; Wu et al., 2016). WUE naturally varies among individuals, both within and among species (Anderegg, 2015; Donovan et al., 2007; Heschel et al., 2002; Tortosa et al., 2016; Yoo et al., 2009) and intraspecific variation in WUE traits can be as great as interspecific variation (Messier et al., 2010). Yield effects resulting from intraspecific WUE trait variation are of considerable agricultural interest (Dutra et al., 2018; Sreeman et al., 2018). Interestingly, it has been shown that the WUE traits of some species of trees alter the photosynthetic parameters and survival of neighboring trees (Bunce et al., 1977), suggesting potential neighbor-scale responses that can dramatically influence the yield of populations. However, few studies have examined the role of WUE trait variation in causing either population-scale or neighbor-scale responses that result in changes in population growth and yield, and this is largely due to the complications that emerge in studying variations in WUE phenotypes.

Studying the scale at which RIs respond to signals that originate from variation in WUE traits requires the anticipation of factors that confound what RIs may be responding to. WUE is calculated as the ratio of carbon assimilation to transpirational water loss, and WUE phenotypes typically result from altered stomatal behavior. However, stomatal phenotypes can also cause pleiotropic effects: an individual’s emission of volatiles could increase as a consequence of increased stomatal conductance, and plant volatiles can change the growth and yield of neighboring plants (Fincheira and Quiroz, 2018; Ninkovic et al., 2016). In addition, the resulting population- or neighbor-scale effects could be due only to changes in transpiration, rather than to changes in WUE. As the frequency of plants with low WUE (high transpiration) increases in a population, the availability of soil water to the population will decrease proportionally (Zea-Cabrera et al., 2006). RIs may change their growth and yield in response to differences in soil water availability, rather than to the abundance of low WUE plants in a population. Therefore, controlling for soil water availability independently of the frequency of plants with different WUE traits is essential to determine the scale at which signals from WUE trait variation alter population growth and yield. Pot experiments conducted in the glasshouse can control for soil water availability but can also confound the analysis of neighbor-scale responses. As planting setups are reduced in size to, for example, the commonly used paired-plants-sharing-a-pot experimental design, individual plants will have more contact with pot sides than with their neighbors (Limpens et al., 2012; Poorter et al., 2012). The ecological relevance of variation in WUE traits is best evaluated in field populations, but standardizing water availability across populations in the field can be a challenge and thus combinations of field and pot experiments are useful for studying the scale at which WUE trait variation causes changes in population growth and yield. With plants that vary in WUE as a result of single-gene manipulations, investigations into the scale (neighbor- or population-scale) at which signals from WUE trait variation are communicated, could be more easily conducted.

Mitogen-activated protein kinases (MAPKs) are part of a conserved signaling cascade essential in eukaryotes. The downstream targets of this phosphorylation cascade, such as transcription factors, enable specific plant responses through changes in plant growth and development (Xu and Zhang, 2015). Mitogen-activated protein kinase 4 (MPK4) in *Nicotiana attenuata* and its homologues MPK12 in *Arabidopsis thaliana* and MPK4/MPK4L in *N. tabacum* have been implicated in responses to herbivore damage (Gomi et al., 2005; Hettenhausen et al., 2013; Yanagawa et al., 2016), bacterial inoculation (Hettenhausen et al., 2012), changes in exogenous and endogenous abscisic acid (ABA) and hydrogen peroxide levels (Des Marais et al., 2014; Hettenhausen et al., 2012; Jammes et al., 2009), vapor pressure deficits (Des Marais et al., 2014) and ozone levels (Gomi et al., 2005; Yanagawa et al., 2016). Most of these responses involve the regulation of stomatal structure and function: silencing *NaMPK4* or *NtMPK4/L* by RNA interference (Na-irMPK4 and Nt-MPK4/L-IR, respectively) or knocking out *AtMPK12* (At-*mpk12*) results in plants with larger stomata and stomatal apertures, and varying disruptions in stomatal closure (Des Marais et al., 2014; Gomi et al., 2005; Hettenhausen et al., 2012; Marten et al., 2008; Yanagawa et al., 2016).

The alteration of stomatal phenotypes by *MPK4/12* strongly influences WUE. Na-irMPK4, Nt-MPK4/L-IR and At-*mpk12* all have increased transpiration rates which can be attributed to increased stomatal conductance (Des Marais et al., 2014; Gomi et al., 2005; Hettenhausen et al., 2012; Yanagawa et al., 2016). For Na-irMPK4 and At-*mpk12*, this increase in transpiration rates has been shown to dwarf the associated increases in assimilation rates, resulting in low WUE (Des Marais et al., 2014; Hettenhausen et al., 2012). However, previous glasshouse studies which tested whether the presence of *MPK4/12*-derived WUE phenotypes results in individual growth and yield effects in paired-plant-in-a-pot interactions, did not control for soil water availability or investigate the effects at different scales within a population (Des Marais et al., 2014; Hettenhausen et al., 2012). To our knowledge, no study has investigated whether variation in the abundance of a low WUE trait, generated from the silencing of a single gene, affects population yield, nor has determined whether this occurs at a neighbor or population scale. Here, we conduct such a study using *N. attenuata* populations.

The wild tobacco *N. attenuata* grows in xeric habitats in the western United States, where water limitation and WUE are selective factors throughout the growing season. *N. attenuata* typically grows in genetically diverse populations, in near-monocultures (Baldwin and Morse, 1994; Baldwin et al., 1994), and is known to respond to genetically different neighbors: a *N. attenuata* accession collected in Utah (UT) sharing a pot with an accession collected in Arizona (AZ) produces significantly smaller stalks than when sharing a pot with another UT plant (Glawe et al., 2003). Additionally, when UT plants are grown in mono-versus mixed-cultures with isogenic lines having genetically modified defense responses, UT plants may suffer less canopy damage from attack by the specialist herbivore *Tupiocoris notatus* (Adam et al., 2018) or more frequent attack by the stem-boring specialist *Trichobaris mucorea* (Schuman et al., 2015). *N. attenuata* plants naturally interact with AMF in the field, establishing networks of connected plants that have the potential to significantly complicate the analysis of these individuals’ responses to neighbors or populations with different WUE phenotypes. AMF are known to change the soil water availability and transport among individuals in populations based on each individual’s ability to interact with AMF (Egerton-Warburton et al., 2007; Reynolds et al., 2003; Yang et al., 2013). Calcium and calmodulin-dependent kinase (CCaMK) is required for successful plant symbiosis with AMF (Lévy et al., 2004), and the abrogation of *CCaMK* expression provides a valuable tool to disconnect plants from AMF networks in the field (Groten et al., 2015). Field plantations of transgenic *N. attenuata* crossed with *CCaMK*-deficient transgenic lines (irCCaMK) in its native habitat, the Great Basin Desert, allow for the study of population growth and yield effects resulting from the varying of traits and the scales at which these may occur. Here, we use single gene manipulations in the background of irCCaMK lines to isolate the complications due to AMF-mediated interactions from responses to variation in WUE traits at the neighbor or population scale.

We investigated the scales at which variation in abundance of low WUE *N. attenuata* plants, created through the abrogation of *MPK4* expression, change individual and population growth and yield. We used a previously characterized irMPK4 line (Hettenhausen et al., 2012) and varied percentages of this line in field populations with empty-vector (EV) control plants, both crossed with irCCaMK. We observed increased yields in populations with low percentages of *MPK4*-deficient plants (‘low-irMPK4’). To exclude soil water availability effects, we grew homozygous irMPK4 and EV lines in glasshouse populations under equal water availability and again observed increased yields in low-irMPK4 populations. The configuration of immediate neighbor genotypes in these populations did not explain changes in individual yields. We further tested responses at the neighbor scale by growing single plants and mono- and mixed-genotype pairs of EV and irMPK4 under conditions of equal water availability. We found no changes in growth or yield between pair types; therefore, neighbor-scale responses appeared unlikely. Between single and paired pots, we found that EV changes its growth and yield in response to neighbors, but irMPK4 does not, indicating that irMPK4 plants have lost their ability to be RIs. We analyzed photosynthetic parameters of individuals in all planting types across glasshouse and field experiments to evaluate if signals interpreted by the EV RIs manifest in changes of photosynthetic parameters. In the glasshouse, changes in EV plants’ photosynthetic parameters alone did not explain the yield increases in low-irMPK4 populations, although EV and irMPK4 plants did simultaneously achieve peak values of WUE in these populations. Importantly however, EV and irMPK4 plants did not differ in WUE phenotypes in field populations, and therefore irMPK4’s WUE phenotype was likely not the cause for the effect at the population scale. We investigated whether the lack of response to neighbors in irMPK4 was root-derived by micro-grafting EV shoots to irMPK4 roots. Silencing *MPK4* in roots was not sufficient to abrogate *N. attenuata’s* response to neighbors in terms of changes in fitness correlates. From these results, we infer a novel function of *MPK4* in mediating *N. attenuata*’s response to neighbors, and in particular shoot *MPK4* in regulating fitness correlate changes in response to neighbors. In addition, *MPK4*-deficient plants at low abundance in population increase total population fitness correlate production likely through information transferred at the population scale, independent of irMPK4’s WUE phenotype.

## Results

### irMPK4xirCCaMK crosses are silenced in *MPK4* and abrogated in AMF associations

*N. attenuata* silenced in the expression of *MPK4* (irMPK4) have a low water-use efficiency (WUE) phenotype in comparison to empty-vector (EV) control plants in the glasshouse (Figure 2). The loss of stomatal control increases transpiration rates to levels that surpass the increases in assimilation rates, consequently decreasing WUE, calculated as the ratio of assimilation:transpiration rates (Hettenhausen et al., 2012).

**Figure 1.**
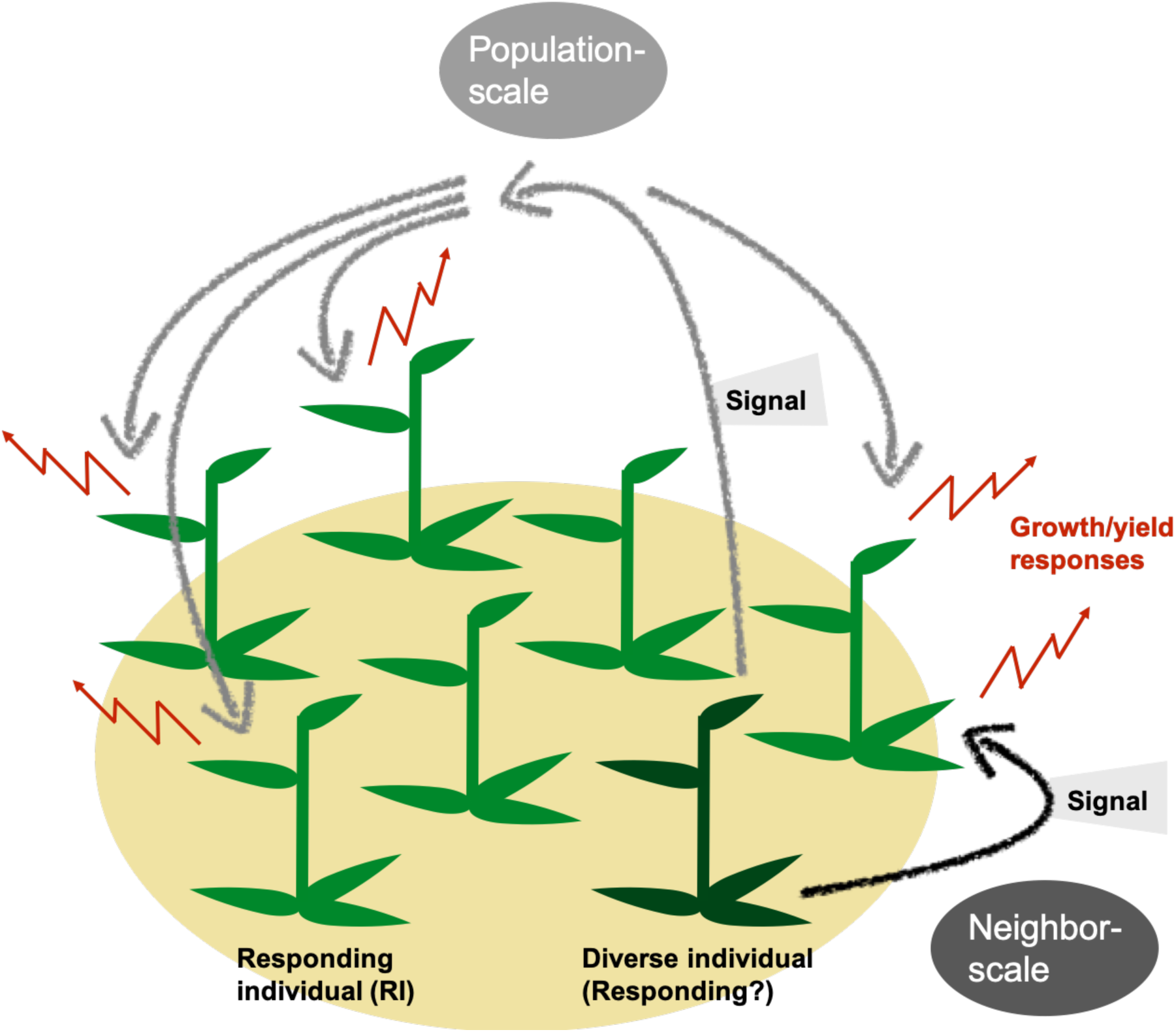
Genetically diverse plants in a population may produce signals interpreted by other plants at the neighbor- or population-scale, changing growth and yield of individuals and populations. Genetic variation in populations can change population growth and yield through changes in individual growth and yield either locally within the population (neighbor-scale) or across the population (population-scale). At the neighbor-scale (black), a plant may produce a signal which only reaches its immediate neighbors, of which some may respond to the signal (responding individuals, RIs). RIs responses may include changes in growth and yield which cumulatively change a population’s growth and yield (red). At the population-scale (gray arrows), a rare plant (dark green) may produce a signal which can be received across the population by RIs.

**Figure 2.**
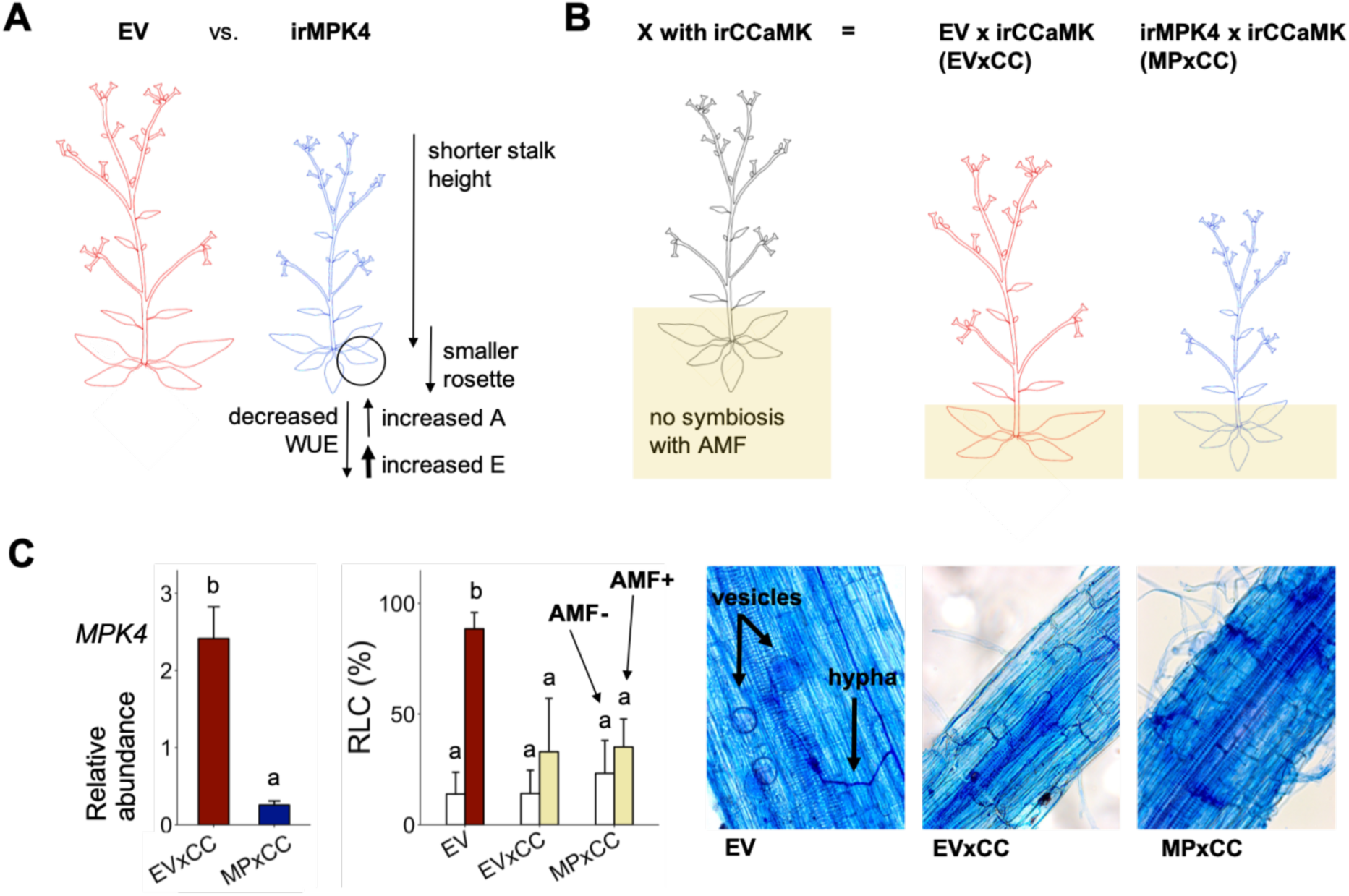
Characterization of EV, irMPK4, EVxirCCaMK and irMPK4xirCCaMK plants. **(A)** irMPK4 *Nicotiana attenuata* plants, silenced in *mitogen-activated protein kinase 4*, have disrupted stomatal control resulting in increased rates of leaf transpiration (E) which surpass increases in rates of leaf carbon assimilation (A) and therefore decrease water-use efficiency (WUE) in comparison to empty-vector (EV) plants. irMPK4 stalks and rosettes are smaller than those of EV. **(B)** irCCaMK *N. attenuata* plants, silenced in *calcium and calmodulin-dependent protein kinase*, are not able to associate with arbuscular mycorrhizal fungi (AMF). These were crossed with EV and irMPK4 to create EVxirCCaMK (EVxCC) and irMPK4xirCCaMK (MPxCC) lines hemizygous for each of the transgenes. **(C)** The reduction of *mitogen-activated protein kinase 4* relative transcript abundances (*MPK4*) in hemizygous MPxCC plants relative to EVxCC (left panel, n = 9 for EV, 13 for irMPK4) is comparable to that of homozygous irMPK4 plants relative to EV (Figure S1, n = 8 for EV/EV and 6 for irMPK4/irMPK4). EVxCC and MPxCC roots which were inoculated with an arbuscular mycorrhizal fungus, *Rhizophagus irregularis* (AMF+), did not show significant increases in comparison to un-inoculated counterparts (AMF-) in root length colonization (RLC, % + CI, n = 7 for EVxCC and 8 for MPxCC), as is the case for EV (n = 8). Images by E.M.: vesicles and hyphae can be seen in trypan blue-stained AMF+ EV roots, but not in AMF+ EVxCC and MPxCC roots.

Populations of plants growing in the field are commonly interconnected by arbuscular mycorrhizal fungal (AMF) networks that are known to influence access to water and nutrients in the plant rhizosphere (Egerton-Warburton et al., 2007; Reynolds et al., 2003; Yang et al., 2013). As silencing the expression of *NaCCaMK* disconnects plants from AMF networks (Groten et al., 2015), we crossed isogenetic, homozygous irCCaMK plants with homozygous EV and irMPK4 lines to generate hemizygous EVxirCCaMK (EVxCC) and irMPK4xirCCaMK (MPxCC) lines (Figure 2B), which were used for field experiments. The hemizygous crosses retained *MPK4* silencing of the homozygous irMPK4 lines: MPxCC showed an 87% reduction of *MPK4* transcript accumulation relative to EVxCC in the field (Figure 2C), whereas irMPK4 had 83% silencing efficiency relative to EV in the glasshouse (Figure S1). To evaluate the abrogation of AMF associations under controlled conditions, we grew the EVxCC and MPxCC crosses in the glasshouse with and without live AMF inoculum (*Rhizophagus irregularis*). While EV was highly colonized in comparison to a non-inoculated control (Figure 2C, LM, *emmeans*_(EV, AMF-[N=8] to AMF+[N=8])_, t = −8.894, p = < 0.0001), both EVxCC (*emmeans*_(EVxCC, AMF-[N=7] to AMF+[N=7])_, t = −2.105, p = 0.4251) and MPxCC (*emmeans*_(MPxCC, AMF-[N=8] to AMF+[N=8])_, t = −1.417, p = 0.8453) did not differ in root length colonization (RLC) with or without AMF. Trypan blue-staining of roots showed the establishment of vesicles and hyphae in EV, but not in EVxCC and MPxCC plants (Figure 2C). From these results, we conclude that the hemizygous crosses retain their *MPK4* silencing and do not associate with AMF.

### Field populations with low percentages of MPxCC plants have increased yield

In order to evaluate if the percentage of *MPK4*-deficient plants influences population yield under field conditions, growth and yield of EVxCC and MPxCC individuals in populations with varying percentages of MPxCCs (Figure 3A; Figure S2) were measured, and population additive totals (PATs) were estimated (see *Plant growth and yield measurements*).

**Figure 3.**
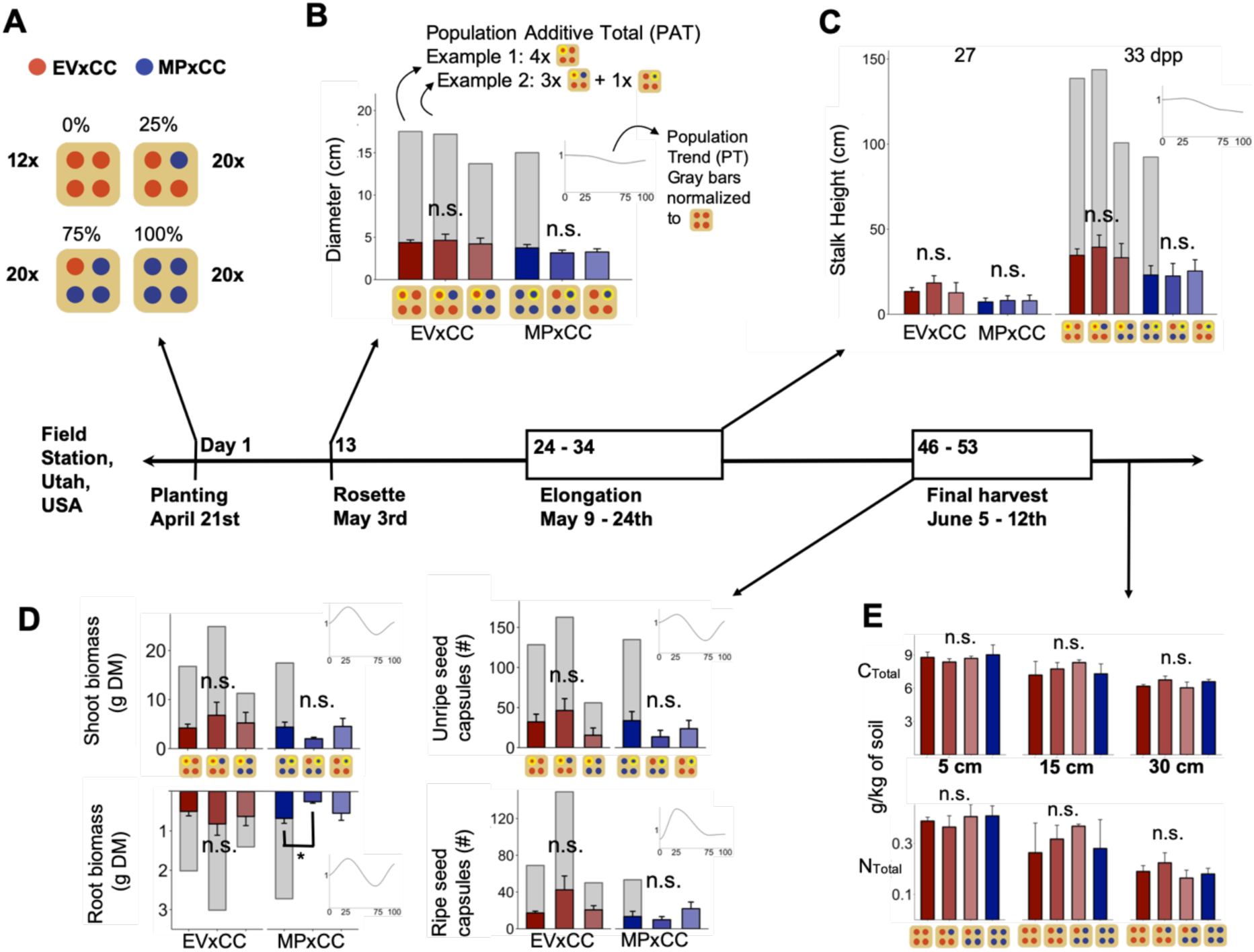
In the field, populations with low percentages of irMPK4 crosses show trends towards increased biomass and reproductive output. **(A)** Field populations of four plants around a central water dripper were planted with varying percentages of irMPK4 crosses: four EVxCC (0%, n = 12), three EVxCC and one MPxCC (25%, n = 20), one EVxCC and three MPxCC (75%, n = 20), or four MPxCC plants (100%, n = 20; Figure S2). Panels B-D: measurements from individual plants in varying population types (x-axis label indicates the measured plant in population in yellow) are represented by colored bars. Population additive totals (PATs) were estimated by taking the individual means from specific populations and multiplying them by the number of that genotype in each population type (example in B). Insets B-D: PATs were divided by the value of the EVxCC monocultures’ PAT, resulting in relative population totals (PTs, y-axis) which were graphed across irMPK4 percentages in population (x-axis). **(B)** Rosette diameters (cm + CI, n = 11-35) at 13 days post planting (dpp) of individual EVxCC (red) and MPxCC (blue) plants in each population type, with estimated PATs (grey bars) and PT (inset). **(C)** Stalk heights (cm + CI, n = 6-29) at 23, 27 and 33 dpp of individual EVxCC (red) and MPxCC (blue) plants in each population type, with estimated PATs (grey bars) and PT (inset). **(D)** Left panels: shoot biomass (top, grams of dry mass, DM + CI, n = 10-31) and root biomass (bottom, g DM + CI, n = 8-31) harvested between 46 and 53 dpp of individual EVxCC (red) and MPxCC (blue) plants from each population type, with estimated PATs (grey bars) and PT (inset). Right panels: unripe seed capsules (top, count + CI, n = 4-14) and ripe seed capsules (bottom, count + CI, n = 3-17) collected between 46 and 53 dpp of individual EVxCC (red) and MPxCC (blue) from each population type, with estimated PATs (grey bars) and PT (inset). **(E)** Soil cores were taken from below the center dripper of each population type at the end of the season (54 dpp, 5 to 30 cm below ground level, n = 3) and homogenized portions of these cores were analyzed (mg/kg + CI) for total carbon (C_total_) and nitrogen (N_total_) - displayed here - and organic carbon (C_organic_), inorganic carbon (C_inorganic_), copper (Cu), iron (Fe), potassium (K), phosphorus (P), and zinc (Zn, Figure S3).

At the individual level, rosette diameters (Figure 3B) and stalk heights of EV and irMPK4 plants measured throughout the season did not differ significantly when compared across the different population types. The biomass and fitness correlate measurements of these individuals at the end of the season also did not differ within either genotype across populations (Figure 3D). Only root biomass was significantly reduced in MPxCC plants in populations with 75% MPxCC compared with 100% MPxCC populations (Figure 3D, LMER, *emmeans*_(MPxCC, 75%[N=4] to 100%[N=3])_, t = 3.243, p = 0.0179).

At the population level, rosette and stalk height PATs and the representations of these PATs as normalized line graphs (population trend, PT), showed additive patterns, in which the calculated population parameters decrease linearly from EVxCC to MPxCC monocultures (insets, Figure 3B, 3C). The biomass and fitness correlate PATs and PTs showed a sinusoidal, non-additive pattern, in which shoot biomass, root biomass, unripe seed capsules, and ripe seed capsules at the population level attained maximum values in populations with 25% MPxCC (‘low-irMPK4’) and minimum values in 75% MPxCC (‘high irMPK4’) populations (Figure 3D).

Plants with low WUE are thought to increase the flow of water-soluble nutrients to the immediate area around their roots as a consequence of excessive transpiration rates (del Amor and Marcelis, 2005; Zea-Cabrera et al., 2006). Therefore, we collected soil cores at 5, 15 and 30 cm below the center of each population type and quantified total carbon (C_total_), nitrogen (N), inorganic carbon (C_inorg_), organic carbon (C_org_), copper (Cu), iron (Fe), potassium (K), phosphorus (P) and zinc (Zn) concentration (Figure 3E; Figure S3). At each depth, there were no significant differences among populations for any nutrient except for C_inorg_ which was slightly increased at 5 cm depth in the 0% populations (EVxCC monoculture), at 15 cm in 75% MPxCC populations, and at 30 cm in 100% MPxCC populations (Figure S3A). We observed no increases at any soil depth in the 25% MPxCC populations (Figure 3E), where the PAT and PT maximums occurred (Figure 3D). Furthermore, the percentage of MPxCC plants in populations did not significantly predict soil moisture at any sampling depth (Figure S4A; R^2^ = 0.527, F_(15, 44)_ = 5.374, p = 6.097e-06). From these results we conclude that increasing the percentage of MPxCC plants in populations under field conditions leads to a non-additive trend in population yield, unrelated to soil nutrient and moisture availability, with maximum population yields occurring in low-irMPK4 populations.

### Glasshouse populations with low percentages of irMPK4 plants and equal water availability have increased yield

To evaluate whether the water-use phenotype of irMPK4 plants contributed to differences in water availability for populations in ways that were undetectable in the field, we created populations with increasing percentages of irMPK4 (0, 17, 50, 83 and 100%) in the glasshouse (Figure 4A, bottom; Figure S5A) and controlled for water availability among populations (Figure S5B; see *Water treatments* in the Methods section).

**Figure 4.**
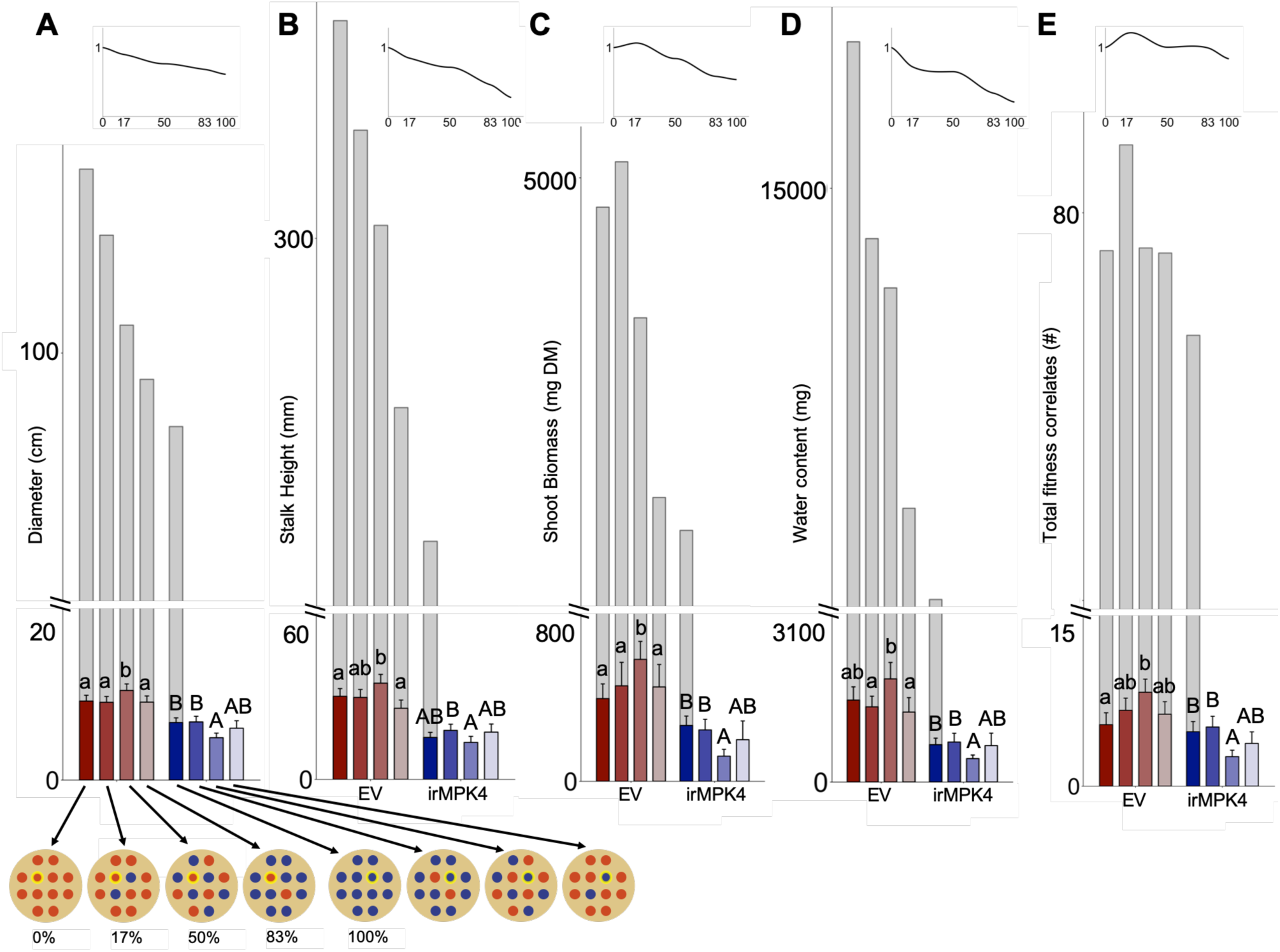
In the glasshouse under equal water availability, populations with low percentages of irMPK4 plants show trends towards increased biomass and reproductive output. **(A)** Glasshouse populations consisted of 12 plants with varying percentages of irMPK4 plants (first five populations, x-axis of A): 12 EVs (0%), 10 EVs and 2 irMPK4s, 6 EVs and 6 irMPK4s (50%), 2 EVs and 10 irMPK4s (83%), or 12 irMPK4s (100%; Figure S5). Each population was watered in proportion to its daily consumption of water to ensure equal water availability across populations. Measurements were made on plants in the center of the population (yellow, x-axis of A) to minimize the influence of pot sides. Panels A-E: measurements from individual plants in different population types are represented by colored bars. Population additive totals (PATs) were estimated by taking the individual means from specific populations and multiplying them by the number of that genotype in each population type (one grey bars for each population type, displayed at their first population type occurrence on the x-axis). Insets B-D: PATs were divided by the value of the EV monocultures’ PAT, resulting in relative population totals (PTs, y-axis) which were graphed across irMPK4 percentages in population (x-axis). **(B)** Rosette diameters (cm + CI, n = 11-35) of EV (red) and irMPK4 (blue) individuals in each population type at 30 days post potting (dpp), with estimated PATs (grey bars) and PT (inset). **(C)** Stalk heights (mm + CI, n = 21-41) of EV (red) and irMPK4 (blue) individuals in each population type at 30 dpp, with estimated PATs (grey bars) and PT (inset). **(D)** Shoot biomass (mg + CI, n = 21-41) of EV (red) and irMPK4 (blue) individuals in each population type at 50 dpp, with estimated PATs (grey bars) and PT (inset). **(E)** Water content (mg + CI, n = 22-41), calculated as the difference between fresh and dry shoot biomass, of EV (red) and irMPK4 (blue) individuals in each population type at 50 dpp, with estimated PATs (grey bars) and PT (inset). **(F)** Total fitness correlates (buds, flowers, unripe and ripe seed capsules; counts + CI, n = 19-44) of EV (red) and irMPK4 (blue) individuals in each population type at 50 dpp, with estimated PATs (grey bars) and PT (inset).

At the individual level, EV plants showed significantly higher growth and yield in 50% irMPK4 populations in comparison to EV plants grown in 0% irMPK4 populations (Figure 4A-E; LME, EV: 0% [n = 40-44] to 50% [n = 21-22], *emmeans*_(Diameter)_, t = −2.796, p = 0.0268; *emmeans*_(Stalk Height)_, t = −2.644, p = 0.0438; *emmeans*_(Shoot Biomass)_, t = −3.859, p = 0.0009; *emmeans*_(Total Fitness Correlates)_, t = −3.825, p = 0.0010). In contrast to the EV plants, irMPK4 plants had significantly smaller rosettes, water contents, and yields in the 50% populations than in 100% irMPK4 monocultures (Figure 4A-E; LME, irMPK4: 100% [n = 31-32] to 50% [n = 22 for all], *emmeans*_(Diameter)_, t = 3.898, p = 0.0006; *emmeans*_(Shoot Biomass)_, t = 5.359, p = <.0001; *emmeans*_(Water Content)_, t = 3.819, p = 0.0010; *emmeans*_(Total Fitness Correlates)_, t = 3.794, p = 0.0011). These changes in EV and irMPK4 individual growth and yield were not correlated with the composition of immediate neighbors to the measured plant: for example, EV plants in 50% and 83% irMPK4 populations had four immediate irMPK4 neighbors, and yet their growth and yields were not equivalent.

At the population level, rosette diameter, stalk height, and water content PATs and PTs were additive (grey bars and insets, Figure 4A, B, D). However, shoot biomass and fitness correlate PATs and PTs were non-additive, with both measurements reaching maximum values in populations with 17% irMPK4 (grey bars and insets, Figure 4C, E). From these results, we conclude that low-irMPK4 populations produce maximum yield both in the glasshouse and field, and these differences are independent of potential differences in population water availability resulting from irMPK4’s WUE phenotype. These results also appear independent from immediate neighbor configurations, indicating that involved responses likely occur at the population rather than the neighbor scale.

### *MPK4* is necessary for *N. attenuata*’s growth and yield responses to neighbors

To test if the increased yields in low-irMPK4 glasshouse (Figure 4E) and field (Figure 3D) populations results from a signal generated by irMPK4 presence at the neighbor scale, we investigated the growth and yield of EV and irMPK4 in monoculture or mixed pairs, in comparison to plants planted alone under equal water availability conditions (Figure 5A). EV plants with an EV (Mono) or irMPK4 (Mix) neighbor had smaller rosettes (Figure 5B, LM, EV: *emmeans*_(Single[N=12] to EV Mono[N=24])_, t = 5.131, p = <.0001; *emmeans*_(Single[N=12] to Mix[N=12])_, t = 4.979, p = <.0001) and fitness correlate production (Figure 5F, LM, EV: *emmeans*_(Single[N=11] to EV Mono[N=22])_, t = 2.434, p = 0.0447; *emmeans*_(Single[N=11] to Mix[N=11])_, t = 3.322, p = 0.0038) than when planted alone (Single). However, this reduction was independent of the neighbor’s genotype (Figure 5B). Shoot biomass was only significantly reduced in EV plants with an irMPK4 neighbor in comparison to individually-grown EV plants (left panel, Figure 5D, LM, EV: *emmeans*_(Single[N=11] to EV Mono[N=22])_, t = 2.434, p = 0.1571; *emmeans*_(Single[N=11] to Mix[N=10])_, t = 3.322, p = 0.0164). EV plants did not have different water contents when planted alone or in pairs (left panel, Figure 5E). irMPK4 plants showed no differences in their rosette growth, water content, or yield when planted in pairs as compared to being grown alone (Figure 5B; Figure 5D-F).

**Figure 5.**
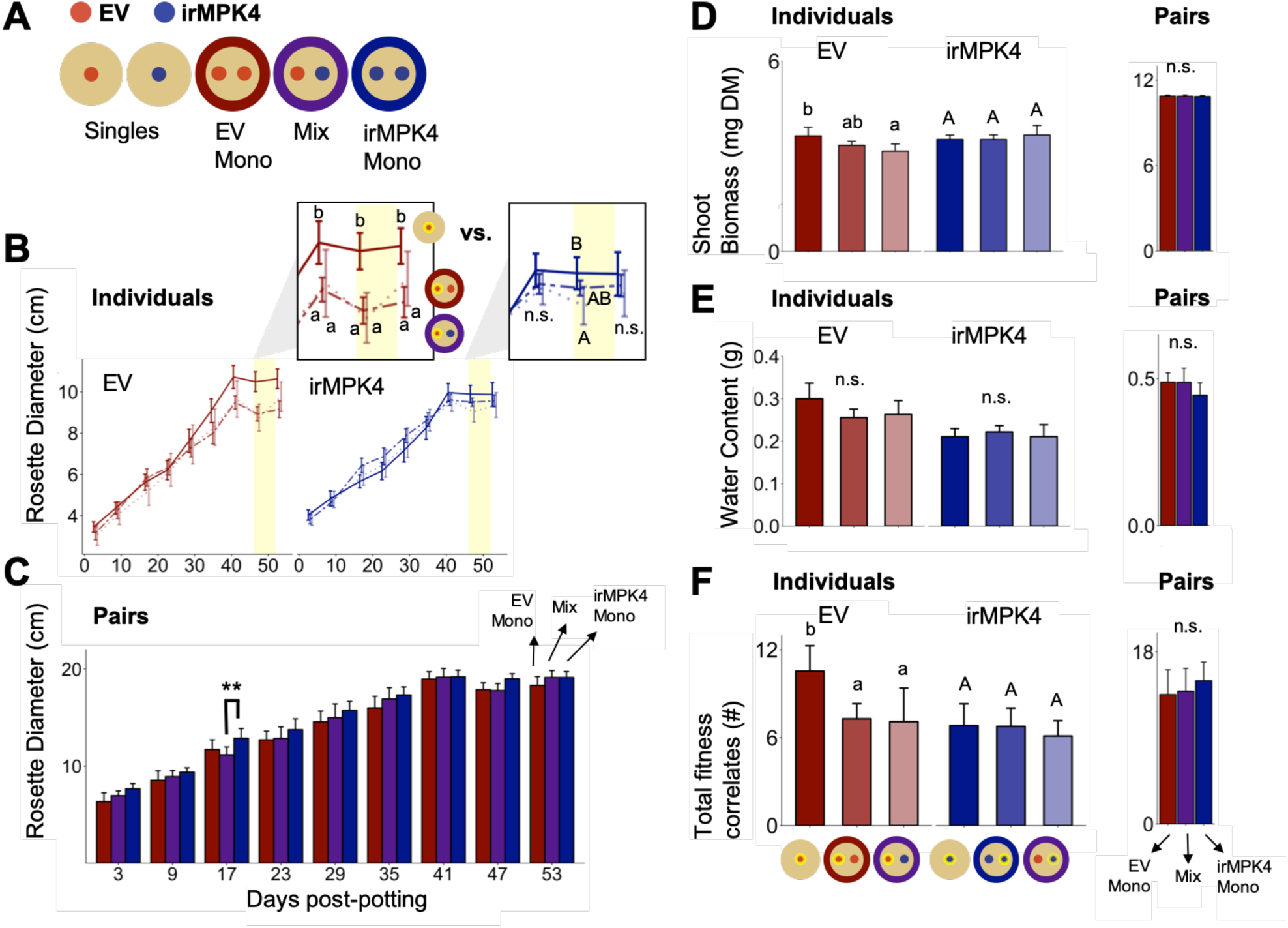
In the glasshouse under equal water availability, EV but not irMPK4 plants have reduced growth in the presence of a neighbor. **(A)** EV and irMPK4 were planted either as singles or in mono- or mixed-culture pairs. All pots were watered based on daily individual consumption to ensure equal water availability. **(B)** Rosette diameters (cm ± CI, n = 11-24) of EV (red) and irMPK4 (blue) individuals in each pot type (Single: solid line; Mono: dashed line; Mix: dotted line), from 3 to 53 days post potting (dpp). **(C)** Cumulative rosette diameters (cm ± CI, n = 11-12) from each pot type (EV Mono: red; Mix: purple; irMPK4 Mono: blue) from 3 to 53 dpp. **(D)** Left panel: shoot biomass (g of dry mass, DM + CI, n = 10-22) of EV (red) and irMPK4 (blue) individuals in each pot type at 71 dpp. Right panel: cumulative shoot biomass (g ± CI, n = 9-11) from each pot type at 71 dpp. **(E)** Left panel: water content (g + CI, n = 8-22), calculated as the difference in grams between fresh and dry shoot biomass, of EV (red) and irMPK4 (blue) individuals in each pot type at 71 dpp. Right panel: cumulative water content (g ± CI, n = 7-11) from each pot type at 71 dpp. **(F)** Left panel: total fitness correlates (buds, flowers, unripe and ripe seed capsules; count #s + CI, n = 9-22) of EV (red) and irMPK4 (blue) individuals in each pot type at 71 dpp. Right panel: cumulative total fitness correlates (count # ± CI, n = 9-10) from each pot type at 71 dpp.

Rosette diameters, shoot biomass, water contents, and fitness correlates of individuals in each type of pair were summed for total outputs per pair type, but no differences were found among pairs for any measurement (Figure 5C; right panels, Figure 5D-F). From these results we conclude that *MPK4* is required for *N. attenuata* growth and yield responses to a neighbor. However, neighbor-scale yield responses of EV and irMPK4 plants in pairs did not have consequences for the total growth and yield of pairs.

### EV and irMPK4 achieve peak WUE in low-irMPK4 populations in the glasshouse, but photosynthetic parameters do not differ in the field

To determine if the WUE phenotypes of EV and irMPK4 plants in glasshouse or field populations change with the percentage of *MPK4*-deficient plants, potentially causing neighbor- or population-scale effects that increase yield (Figure 1), we measured leaf photosynthetic parameters (assimilation rate, transpiration rate, stomatal conductance) and calculated the WUE of all individuals in both glasshouse and field experiments.

In the glasshouse paired experiment, all measured leaf photosynthetic parameters of EV and irMPK4 plants in single pots were as previously reported (Figure 2A), with irMPK4 plants having significantly higher assimilation rates, transpiration rates, and stomatal conductance than EV plants, and significantly lower WUE (Figure 6A; Assimilation, LM, EV_Single[N=4]_-irMPK4_Single[N=3]_, t = − 3.947, p = 0.0070; Transpiration, GLS, EV_Single[N=4]_-irMPK4_Single[N=4]_, t = −8.089, p = <.0001; Stomatal conductance, LM, EV_Single[N=3]_-irMPK4_Single[N=4]_, t = −10.147, p = <.0001; WUE, GLS, EV_Single[N=4]_-irMPK4_Single[N=4]_, t = 6.621, p = <.0001). When planted in pairs, EV and irMPK4 plants assimilation rates, transpiration rates, and stomatal conductance were not significantly different in monoculture (Figure 6A, red and blue shadings) versus in mixed culture (Figure 6A, purple shading). EV plants had significantly lower WUE in mixed versus monoculture (Figure 6A; GLS, EV: *emmeans*_Mono[N=6|-Mix[N=4]_, t = 3.723, p = 0.0205), whereas irMPK4 plants showed no significant change in WUE across pair planting types.

**Figure 6.**
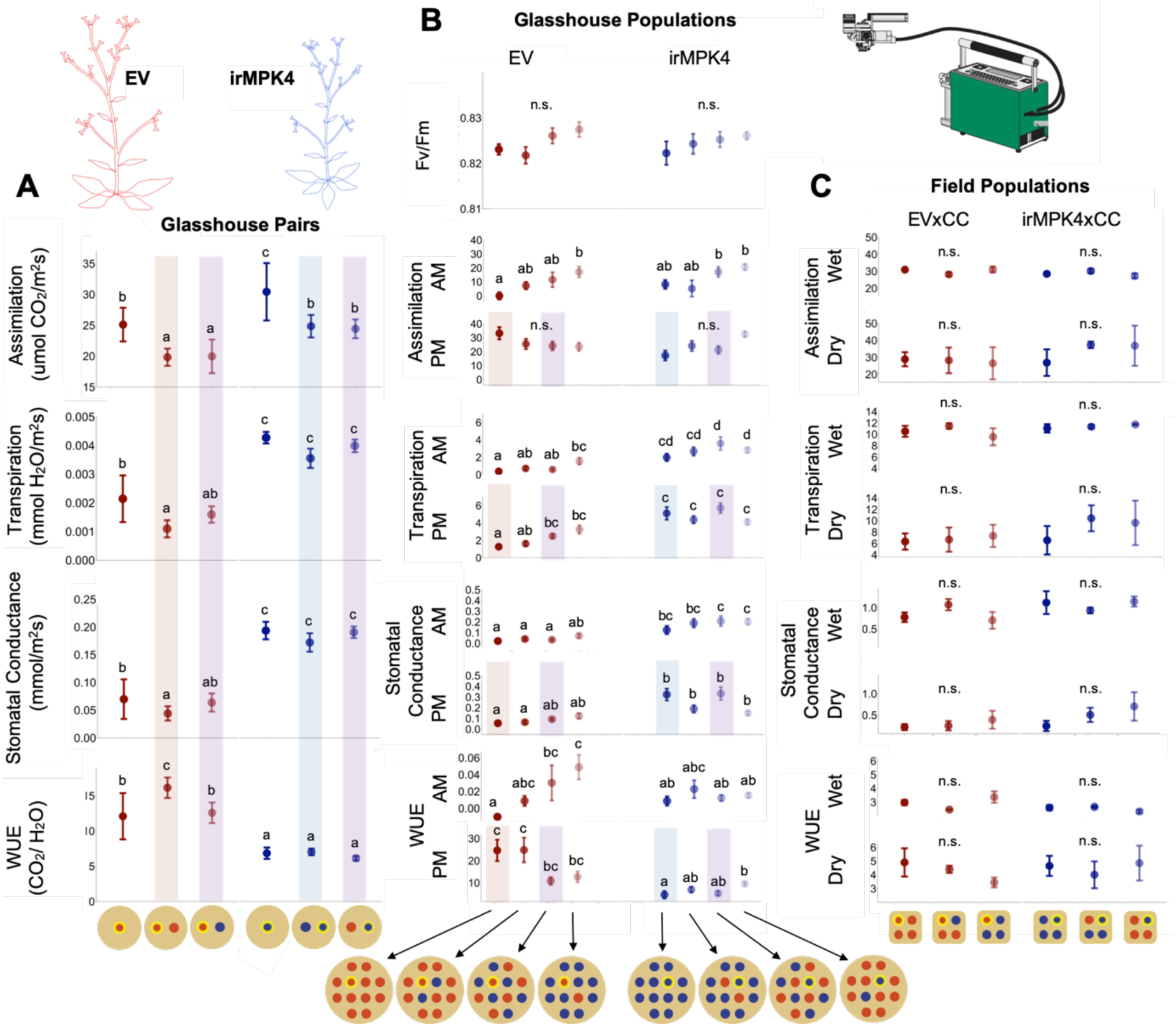
irMPK4’s photosynthetic phenotype is consistent in the glasshouse but not present in the field. **(A)** Assimilation rate, transpiration rate, stomatal conductance and water-use efficiency (WUE; ± CI for all, n = 3-8) of EV (red) and irMPK4 (blue) individuals from each planting type in the paired glasshouse experiment (Figure 5) at 48 days post potting (dpp). For ease of comparison to the population glasshouse experiment, EV’s in 0% irMPK4 populations (EV Mono, red), irMPK4’s in 100% irMPK4 populations (irMPK4 Mono, blue) and both genotypes in 50% irMPK4 populations (Mix, purple) are highlighted. **(B)** Chlorophyll fluorescence, assimilation rate, transpiration rate, stomatal conductance and WUE (± SE for all, for clarity in visualization, n = 11-32) of EV (red) and irMPK4 (blue) individuals from each planting type in the population glasshouse experiment at 32 dpp. Measurements include both pre-dawn dark-adapted measurements (AM, 4:00 to 6:00) when plant turgor pressure typically peaks, and noon measurements (PM, 12:00 to 14:00) when plant leaf water resources are in high demand. For comparison to the paired glasshouse experiment, EV’s and irMPK4’s in 0%, 50% and 100% irMPK4 populations are highlighted. **(C)** Assimilation rate, transpiration rate, stomatal conductance and WUE (± SE for all, for clarity in visualization, n = 3) of EV (red) and irMPK4 (blue) individuals from each planting type in the population field experiment at 34 dpp. Measurements include both measurements on recently watered plants (Wet, consisting of plants which received daily watering), and water deficit measurements (Dry, consisting of plants which were not watered for two weeks).

In the glasshouse population experiment, plants were measured at two timepoints: pre-dawn (AM) and noon (PM, see *Gas exchange measurements and water-use efficiency calculations*). The AM timepoint permitted the measurement of dark-adapted chlorophyll fluorescence, which showed no significant differences among EV or irMPK4 individuals in different population types (Figure 6B). The PM timepoint measurements (Figure 6B) were comparable to all measurements in the other glasshouse and field experiments (Figure 6A, C). The PM photosynthetic parameters of the population and paired glasshouse experiments are highlighted to show the consistency of changes in EV individuals in 0% and 50% irMPK4 populations and irMPK4 individuals in 100% and 50% populations to their respective counterparts in pairs with the same percentage of irMPK4 (Figure 6A, B). irMPK4 plants in 100% (blue shading) versus 50% (purple shading) irMPK4 populations showed no significant differences in any photosynthetic parameter, which was consistent with the glasshouse paired experiment (Figure 6B). EV plants in 0% (red shading) versus 50% (purple shading) irMPK4 populations were not significantly different from each other in any parameter except for significantly higher transpiration rates in EV plants in 50% irMPK4 populations compared with those in 0% irMPK4 populations (Figure 6B; LMER, EV: *emmeans*_0%[N=32]-50%[N=16]_, t = −3.744, p = 0.0082). This result, and the lack of significant differences in WUE of EV plants in 0% versus 50% irMPK4 populations, differed from the results of the paired experiment, and indicated that EV plants’ photosynthetic parameters may be affected by the immediate neighbor genotypes rather than solely by the percentage of irMPK4 in a glasshouse population.

In the glasshouse population experiment, both EV and irMPK4 achieved peak WUE (PM measurement) in low-irMPK4 populations (Figure 6B, EV: second data point from the left; irMPK4: last data point from the left), where increased growth and yield were previously observed (Figure 4C, E). However, while the WUE of irMPK4 plants was significantly increased in the low-irMPK4 population in comparison to irMPK4 monocultures (100% irMPK4 populations), the peak WUE of EV plants was not significantly different from that of EV plants in monoculture (0% irMPK4 populations; Figure 6B; LMER, EV: *emmeans*_0%[N=25]-17%[N=16]_, t = −0.521, p = 1.000; irMPK4: *emmeans*_100%[N=32]-17%[N=11]_, t = −3.479, p = 0.0231).

No photosynthetic parameter for either genotype differed significantly under field conditions, whether in an irrigated (Wet) or non-irrigated (Dry, see *Water treatments* in Materials and Methods) part of the field plot (Figure 6C). Therefore, we conclude that the WUE phenotype of irMPK4 is not likely to have accounted for the greater growth and yield of plants in low-irMPK4 populations.

### Shoot *MPK4* is required for *N. attenuata* to alter its fitness correlate production in response to a neighbor

To test the tissue-specific function of MPK4 expression in plant yield responses to a neighbor, we created chimeric plants by micro-grafting irMPK4 roots to EV shoots (heterografts), EV roots to EV shoots (EV homografts) and irMPK4 roots to irMPK4 shoots (irMPK4 homografts; Figure 7A). We grew the grafts under conditions of equal water availability, with or without an ungrafted EV neighbor. Photosynthetic parameter profiling of these grafts revealed that the heterografts were similar to EV homografts in assimilation, transpiration, stomatal conductance, and WUE within planting type (Figure 7B), while the irMPK4 homografts showed significantly higher transpiration rates and stomatal conductance and lower WUE (Figure 7B, Table 1). Hetero- and homo-irMPK4 grafts retained similar levels of *MPK4* silencing in roots or roots and shoots, respectively (Figure S1).

**Figure 7.**
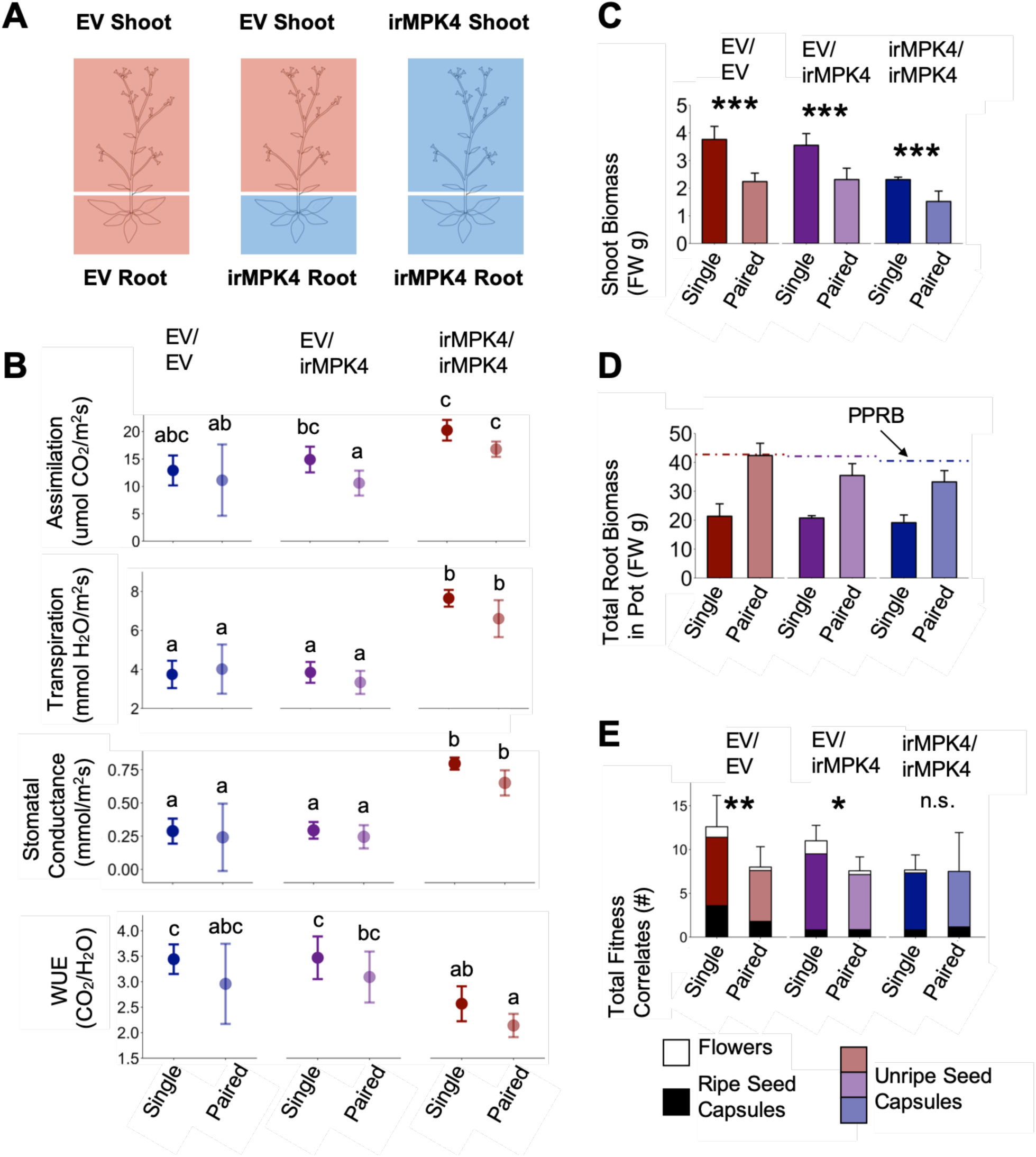
In the glasshouse under equal water availability, shoot expression of *MPK4* mediates changes in *N. attenuata* reproductive output in response to neighbors. **(A)** irMPK4 roots were micro-grafted onto EV shoots, producing plants deficient in *MPK4* in the root but not in the shoot (EV/irMPK4, Figure S1). These were compared to EV/EV and irMPK4/irMPK4 grafts as controls. All three plant types were grown both as singles and in pairs with an ungrafted EV neighbor, under conditions of equal water availability. **(B)** EV/irMPK4 have photosynthetic parameters comparable to those of EV/EV as shown in the assimilation rates, transpiration rates, stomatal conductance and water-use efficiency (WUE; ± CI for all, n = 3-7) of single and paired plants of each grafting type (EV/EV: red; EV/irMPK4: purple; irMPK4/irMPK4: red). **(C)** Shoot biomass (grams of fresh mass, g FM + CI, n = 4-6) of EV (red) and irMPK4 (blue) individuals in each potting type at 50 dpp. **(D)** Root biomass of all plants in each pot (g FM + CI, n = 4-6) for each potting type at 50 dpp. Dashed lines indicate the predicted pot root biomass (PPRB) of the each paired pot, calculated from the addition of means of the individual plants present in the paired pot (EV/EV mean used to represent the EV neighbor). **(E)** Total fitness correlates (flowers, unripe and ripe seed capsules; count #s + CI, n = 5-7) of EV (red) and irMPK4 (blue) individuals in each pot type at 50 dpp. Statistics were calculated on the totals only, although each bar is dissected into its contributing parts: flowers (white), unripe seed capsules (color), and ripe seed capsules (black).

**Table 1.**
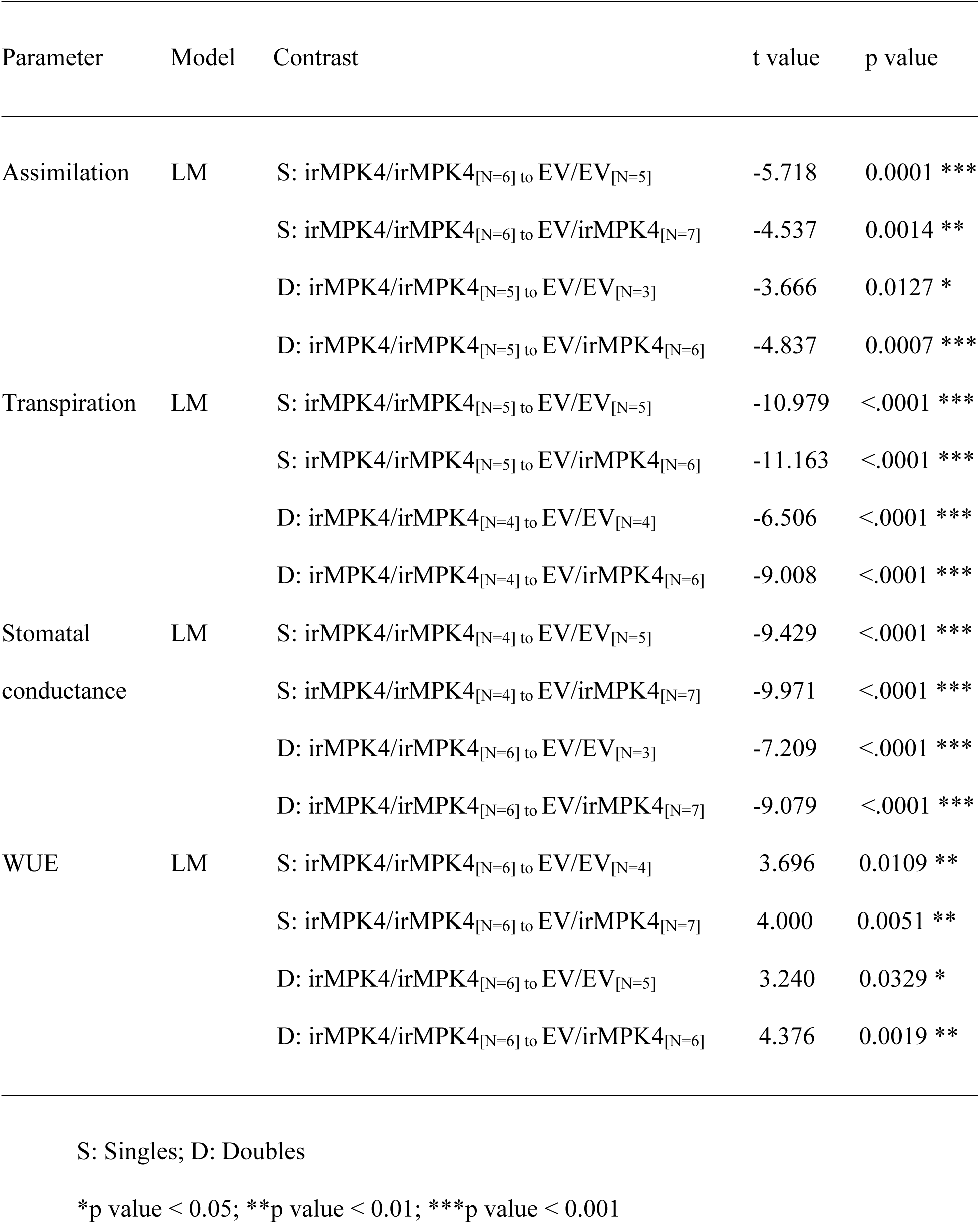
Statistical *emmeans* contrasts within planting treatments for Figure 7B.

All graft types had shoot biomasses that were significantly reduced when plants were grown in pairs versus when planted alone (left panel, Figure 7C; LM, EV/EV: *emmeans*_(Single[N=5] to Paired[N=5])_, t = −7.823, p = <.0001; EV/irMPK4: *emmeans*_(Single[N=4] to Paired[N=6])_, t = −6.232, p = <.0001; irMPK4/irMPK4: *emmeans*_(Single[N=6] to Paired[N=6])_, t = −4.442, p = 0.0001). For root biomasses, the total root biomass per paired pot, representing the roots of two plants, was compared to a predicted pot root biomass (PPRB), calculated by adding the single pot root biomass means of the genotypes present in the paired pot (PPRB, see *Plant growth and yield*). Interestingly, pot root biomasses of EV homograft pairs were equal to their PPRB, while the paired heterografts and irMPK4 homografts had smaller pot root biomasses than estimated by their PPRB (Figure 7D).

Between single and paired plants, irMPK4 homografts did not show a significant difference in fitness correlates (Figure 7E; LM, irMPK4/irMPK4: *emmeans*_(Single[N=13] to Paired[N=10])_, t =0.024, p =0.9813), unlike both the EV homografts and heterografts, which displayed significant reductions in fitness correlates in response to an EV neighbor (Figure 7E, LM, EV/EV: *emmeans*_(Single[N=9] to Paired[N=11])_, t = −2.637, p = 0.0106; EV/irMPK4: *emmeans*_(Single[N=13] to Paired[N=12])_, t = −3.620, p = 0.0006). From these results we conclude that silencing *MPK4* in the roots changes the neighbor-related root biomass production of *N. attenuata*, but *MPK4* in the shoots is required to alter fitness correlate production in response to neighbors.

## Discussion

Individual-level variation in water-use efficiency (WUE, Figure 2A) can change overall population yield (Campitelli et al., 2016; Kenney et al., 2014). This might occur through signals produced by low-WUE plants at several scales within a population’s hierarchical organization (Allen and Starr, 1982): signals might only reach the immediate neighbors (neighbor-scale) and cause them to respond with changes in growth and yield, or the signals may accumulate over the entire population (population-scale) and reach distributed individuals, resulting in some individuals responding (responding individuals, RIs), which have the machinery to do so. Our analyses reveal that low abundances of irMPK4 plants intermixed with EV plants resulted in the highest *N. attenuata* population yields both in the glasshouse and the field (Figures 3D, 4C, 4E). This was not due to differences in soil water availability, which was controlled for in the glasshouse (Figure 4; Figure S5B), nor irMPK4’s WUE phenotype, which was not observed in the field (Figure 6). Interestingly, we find that the signal which was responsible for the yield-increasing responses in low-irMPK4 populations likely occurred at the population scale.

Based on results from previous neighbor-effect studies using low-WUE phenotypes, we initially hypothesized that the neighbor scale (Figure 1, black arrow) would be critical for producing individual changes in yield. In paired-plant competition experiments, the *A. thaliana* CVI mutant (low WUE) produced more seeds than L*er* (high WUE), although the opposite was true when these two lines were planted individually (Campitelli et al., 2016). In addition, the *N. attenuata* irMPK4 line growing in competition with EV caused both genotypes to wilt two days after termination of watering, but when grown individually, only irMPK4 showed this wilting phenotype (Hettenhausen et al., 2012). Despite these changes in the L*er*/EV neighbor responses to plants deficient in *MPK12/MPK4*, in this study we did not see differences among EV plants paired with EV or with irMPK4 in any growth or yield parameters (Figure 5B-F). We considered that paired-plants-in-a-pot experiments may disguise neighbor-scale effects due to a higher proportion of plant roots being in contact with the sides of the pot, rather than with their immediate neighbors. Additionally, the growth and yield responses of plants at the neighbor scale could depend on the identities of several immediate neighbors, as in populations. In our glasshouse population experiments, we only measured plants surrounded by other plants in the population rather than by pot sides (Figure 4; Figure S5A). When comparing the growth and yield results of EV plants among changing numbers of direct irMPK4 neighbors (0 irMPK4s in 0% irMPK4 populations, 2 in 17% populations, and 4 in both 50 and 83% populations), we did not observe a correlation between the two. Given these negative results of our analyses at the neighbor scale, we infer that the responses resulting in observed population yield effect likely occur in relation to a factor at the population scale.

We predicted that irMPK4’s WUE phenotype was likely responsible for the response at the population scale. The low WUE of these plants could change the soil water availability in the population and create a population-scale response to a decreased water table (Zea-Cabrera et al., 2006). This may only have created an increase in RIs growth and yield when the change was minimal (low-irMPK4). Although we did not see differences in soil moisture percentages among different population types in the field (Figure S4), we tested this in the glasshouse by controlling for soil water availability among populations. We hypothesized that without differences in soil water limitation, we would not observe the increases in yield of low-irMPK4 populations, but the increased yield in these populations remained. From these results, we inferred that the yield response was elicited by irMPK4 phenotypes other than its WUE phenotype. Based on previous findings where trees with differing WUE influenced photosynthetic parameters of neighboring trees (Bunce et al., 1977), we examined the photosynthetic parameters of plants in the field and glasshouse. In the field, EV and irMPK4 plants did not differ in their photosynthetic parameters, including in WUE (Figure 6C). In the glasshouse populations, the photosynthetic parameters of EV individuals changed in association with the number of direct neighbors of irMPK4, and EV and irMPK4 plants both reached highest values of WUE in low-irMPK4 populations (Figure 6B). However, we interpreted these responses to be independent of the responses causing the low-irMPK4 population yield increase, given the lack of the irMPK4 WUE phenotype in the field (Figure 6C). We concluded that the population-scale factor responsible for the increased low-irMPK4 population yield originates from an irMPK4 phenotype that is independent of its WUE phenotype.

We considered other types of factors that could alter yields at the population scale. Volatiles can accumulate differently in the headspaces of various plant populations (Schuman et al., 2015) and were shown to be a mechanism by which plants detect and respond to potentially competitive neighbors (Ninkovic et al., 2016; Pierik et al., 2013). Particular volatiles released from microbes can also promote plant growth (Fincheira and Quiroz, 2018). irMPK4 and EV plants are known to have distinct volatile profiles after elicitation by herbivory: in a previous experiment, irMPK4 plants released *trans*-α-bergamotene at 5x the level of EV plants (Hettenhausen et al., 2012). We infer that there may also be a difference in irMPK4 and EV plants’ volatile profiles when not induced, which could be a factor causing responses at the population scale. Further research on the headspace of EV and irMPK4 populations in the field could help to address this question. Alternatively, root exudates can accumulate below-ground in populations and provide identifying information for plants about their neighbors, which may result in plant growth responses (Semchenko et al., 2014). Although the exudate profiles of irMPK4 roots have not been characterized, the results of our paired grafting experiment revealed that silencing *MPK4* only in the root was sufficient to reduce the total root biomass of the irMPK4 plant and of an EV neighbor sharing pot. These reductions were comparable to the reduction in root biomass observed for irMPK4 homografts (silenced in both root and shoot expression) and their EV neighbors (Figure 7D). From these results we infer that irMPK4 in low abundances in populations could reduce root growth through a signal such as root exudates, with the result that EV plant might redirect resources to increase above-ground yield, as is known to occur in the Shade Avoidance Response triggered by phyB responses to far red light (Fragoso et al., 2017; Morelli and Ruberti, 2000). This hypothesis remains to be tested, but perhaps the first step would be identifying the specific tissue responsible for the signal source with experiments conducted with a population of grafted plants.

Our paired grafting experiment revealed tissue-specific requirements for the response of *N. attenuata* plants to neighbor presence: *MPK4* transcripts are required to respond to the presence of a neighbor with altered yield (Figure 5F, 7E), and the presence of these transcripts in the shoot alone are sufficient for this response to occur, as in the EV/irMPK4 heterograft (Figure 7E). We suggest that *MPK4* presence in the shoot could be involved both in the negative regulation information transmitted at the population scale as well in the integration of this signal by RIs. Naturally occurring alleles of *MPK12* in *A. thaliana* have been found to produce varying amounts of MPK12 (Des Marais et al., 2014), indicating that there may yet be unexplored functions of the natural variation of *MPK4/12*, which may relate to its role in mediating a factor that increases population yield. Given that we were unable to investigate the function of shoot-only *MPK4* knockdowns, as the RNAi silencing signals travel from shoots to roots in *N. attenuata* (Fragoso et al., 2011), the use of mutant or natural alleles of *MPK4/12* could allow for the analysis of reciprocal *mpk12/wt* and *wt/mpk12* grafts in *A. thaliana*.

The results of this study are consistent with the well-established result that functionally rare individuals in a population can increase population productivity (Chapin III et al., 1997; Chapin, et al., 1998; Hooper et al., 2005; Naeem et al., 1994; Schulze and Mooney, 1994) and advance our understanding by identifying the scale within populations at which this yield response occurs. Additionally, we identify a single gene, *MPK4,* which when silenced in low abundances in a population is responsible for increasing population yield. We infer that *MPK4* influences population yield through RIs at the population scale by excluding neighbor-scale effects, although future experiments will need to more directly evaluate and identify the population-scale signal. The work contributes to our understanding of how populations may become more productive as a result of greater genetic and functional diversity and suggests that experiments like the ones used in this study which explore the scales at which these effects occur, will identify additional mechanisms that could increase the productivity of agronomic monocultures.

## Materials and Methods

### Plant material and constructs

Characterization of the empty-vector (EV) *Nicotiana attenuata* control line (pSOL3NC, line number A-04-266-3) is described by Bubner and colleagues (Bubner et al., 2006). The irMPK4 line (pRESC5MPK4, line number A-7-163), silenced in the production of MITOGEN-ACTIVATED PROTEIN KINASE 4 (MPK4) through RNAi targeting *MPK4* transcripts, is characterized by Hettenhausen and colleagues (Hettenhausen et al., 2013, 2012). The irCCaMK line (pSOL8CCAMK, line number A-09-1212-1-4), silenced in the production of CALCIUM AND CALMODULIN-DEPENDENT PROTEIN KINASE (CCaMK) through RNAi targeting *CCaMK* transcripts, is characterized by Groten and colleagues (Groten et al., 2015).

EVxirCCaMK (pSOL3NCxpSOL8CCAMK, ‘EVxCC’) and irMPK4xirCCaMK (pRESC5MPK4xpSOL8CCAMK, ‘MPxCC’) crosses were generated by growing EV (second generation, T2), irMPK4 (T2) and irCCaMK (third generation, T3) in the glasshouse and hand pollinating the styles of EV and irMPK4 emasculated flowers with pollen from the anthers of the irCCaMK flowers. Control crosses were made simultaneously: EVxEV (pSOL3NCxpSOL3NC, ‘EVxEV’) and irMPK4xEV (pRESC5MPK4xpSOL3NC, ‘MPxEV’) were pollinated from the anthers of EV flowers. Hand-pollinated flowers were tagged with string, and resulting seed capsules were collected. The ripe seeds from these crosses provided the seed source for the field population experiment (Figure 3). A cross characterization experiment in the glasshouse revealed that EVxEV and EVxCC, as well as MPxEV and MPxCC, were not significantly different in water loss rates per day (Figure S6A, Table S1), or in shoot and root biomass (Figure S6B, C). For all other experiments in the glasshouse, T3 EV and irMPK4 homozygous lines were used.

### Plant growth conditions

Importation and release of transgenic crosses in the field station (Lytle Ranch, Utah, USA) was carried out under Animal and Plant Health Inspection Service (APHIS) import permit numbers 07-341-101n (EV, irMPK4) and 10-004-105m (irCCaMK), and release 16-013-102r.

Glasshouse and field germination and growth were described previously (McGale et al., 2018), with modifications only in planting design. Field plants were planted in four-plant populations in a square design (Figure 3; Figure S2), with 10 cm between each adjacent neighbor. Plants from the glasshouse population experiment were potted in twelve-plant populations (Figure 4; Figure S5), with 5 cm between each adjacent neighbor. Glasshouse plants in both of the paired experiments (grafted and ungrafted) were also planted 5 cm from their neighbor plants.

### Plant growth and yield measurements

For the field experiment (Figure 3), rosette diameter measurements were extracted from photos taken between 19:00 and 20:00, in which each individual plant was pictured next to a standard metal square (5×5 cm) for scale. Plant stalk height measurements were taken as the height from the base of the stalk at the ground level to the highest point of the topmost inflorescence. Plant shoot and root dry biomass were measured by placing respective biological matter in a paper bag inside of a plastic box with ventilation holes of 1 cm diameter drilled through the lid and left to dry for 15 days in the sun, before being removed from the bag and weighed. Unripe seed capsules were counted simultaneously for all plants, immediately before harvesting for shoot and root biomass. Due to APHIS regulations, ripening seed capsules were counted and subsequently removed to prevent opening and releasing seeds into the field; the total ripe capsules collected is presented (Figure 3D).

For all glasshouse experiments, the planting substrate consisted of a bottom layer of large clay aggregate (Lecaton, 816mm diameter, approx. 10% of pot volume), a central layer of small clay aggregate (Lecaton, 24mm diameter, approx. 80% of pot volume) and a top layer of fine sand (approx. 10% of pot volume). This substrate provides optimal drainage in the pots for the purposes of water control, and conditions similar to the sandy, clay nature of the natural habitat of *N. attenuata.* Rosette diameter was measured directly on the plant. Plant stalk height was measured as in the field. Shoot biomass consisted of all above ground matter (severed below the rosette), placed inside a bag for drying at 80°C for 2 h, after which the plant matter was removed from the bag and weighed. The shoot biomass was also weighed for fresh mass, and the water content of the plant at harvest was reported as the difference between the fresh and dry shoot biomasses. All fitness correlates were counted at harvest, including buds (larger than 1mm), flowers (counted as flowers when the corolla became visible by pushing through the sepals), unripe and ripe seed capsules, and the total of all of these together was reported (Figure 4E).

In the grafted experiment, predicted paired root biomass (PPRB) is presented in Figure 7D. This was calculated as the sum of the means of the single graft root biomasses present in the pair (e.g. EV/EV PPRB = 2*(EV/EV single root biomass mean); EV/irMPK4 PPRB = (EV/irMPK4 single root biomass mean) + (EV/EV single root biomass mean)).

### Calculations of population additive totals and population trends

In Figures 3 and 4, data from individuals are represented by red and blue bars. Population additive totals (PAT) are presented as gray bars and population trends (PT) as line graphs.

PATs for each population type were estimated by multiplying the mean of each individual genotype in each type of population by the number of that genotype in that population type. Two example calculations are shown in Figure 3B. Note: in Figure 3, while measurements for six individuals in different population types are shown, only four PATs are presented, as there were only four different population types (i.e. individual bars may originate from the same population type).

To create the PTs, the resulting value of each PAT bar was divided by the PAT of the monoculture EVxCC (or EV, for Figure 4) population and normalized values (y-axis) were plotted against the percentages of *MPK4*-silenced plants in each population type (x-axis).

### Soil moisture and element content

Soil cores were taken from the field by driving a split tube core borer (53 mm, Eijkelkamp, Giesbeek, Netherlands) 30 cm into the ground, and carefully removing it with the core intact. 5 cm pieces of field soil were cut from the core from 0 to 5, 10 to 15, and 25 to 30 cm below ground. Each of these 5 cm thick pieces were weighed, left to dry in the sun in UV-excluding boxes similar to those used for the drying of shoot biomass (see *Plant growth and yield measurements*), and weighed again when dry (determined to be when the mass fluctuated <0.1 g between days). Soil moisture was calculated from each sample (% soil moisture = (fresh soil mass - dry soil mass / fresh soil mass) * 100), taken from 21 to 30 dpp (Figure S4, n = 1 per population).

Soil cores were obtained using the same method at 54 dpp with replication (n = 2-9) to determine the soil content of total, inorganic and organic carbon (C_total_, C_inorg_, C_org_, respectively), nitrogen (N), copper (Cu), iron (Fe), potassium (K), phosphorus (P), and zinc (Zn) in each type of population at the end of the season. These samples were dried at 80°C for 6 h in a drying oven (Figure 3E; Figure S3), and sieved and milled for C_total_ determination (by elemental analyzer), C_inorg_ (by loss-on-ignition), C_org_ (C_total_ - C_inorg_), N (by elemental analyzer) at the Max Planck for Biogeochemistry in Jena, Germany. A separate Cu/Fe/K/P/Zn determination (by microwave digestion via atomic spectroscopy) was also performed.

### Water treatments

Field plant populations were watered every week for 1 h at dusk (20:00 to 21:00) from a central water dripper (2L/h drip rate) present in each population. After 34 dpp, one section of the plot was no longer watered until the final harvest (Figure 3; Figure 6C: “Dry”), while a small subsection was watered two more times (Figure 6C: “Wet”) in order to obtain gas exchange measurements on sections with varying water treatments at 48 dpp. Soil moistures in these two parts of the plot (see *Soil moisture and element content*) were graphed as regression lines from 21-30 dpp to test if results from both of these parts could be summarized together in Figure 3. The part of the plot where soil cores were harvested did not significantly predict soil moisture, nor did the interaction of section with depth or day (Figure S4B, Wet subsection: ‘Part2’; day: ‘variable’; model fit: R^2^ = 0.406, F(7, 147) = 16.04, *p*-value = 1.474e-15), therefore shoot and root biomass, as well as unripe and ripe seed capsule data collected from full populations in both sections were reported together as one mean (Figure 3).

In the glasshouse, all populations and pairs (grafted and ungrafted) underwent the following regimented watering to control for water availability: after potting, they were given establishment watering (soil moisture maintained around 20%) for three weeks, allowing root development to the bottom of the pot for a transition from top watering to bottom watering. After three weeks, pots began to show detectable differences in water loss by population type and consumption-based watering began. For the population and homozygous pair experiment, pots were individually watered daily to a two-day water supply, calculated as:

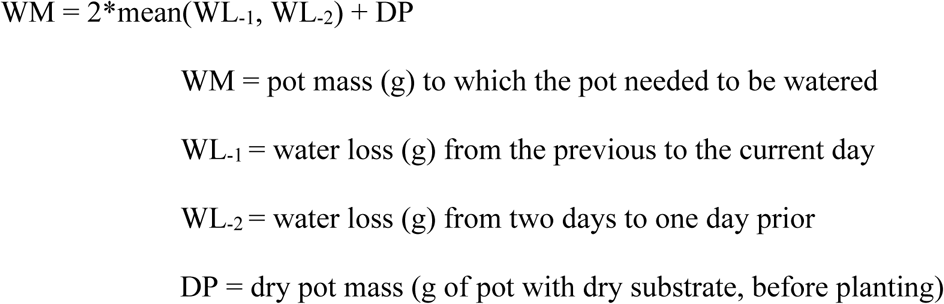

The two-day water supply was chosen as it reflected ecologically-relevant soil moisture at the field site in the natural habitat of *N. attenuata* (Valim et al., 2019). To allow larger growth and thus accentuate growth differences in plants in the grafted pair experiment, the water supply was raised to five days (WM = 5*mean(WL_-1_, WL_-2_) + DP), bringing soil moisture percentages up to 20-30%. The higher soil moisture did not affect the differences in photosynthetic parameters of EV and irMPK4 homozygous grafts compared to those reported for the homozygous EV and irMPK4 plants in the paired experiment (Figure 6A, 7B). There was no significant correlation between the amount of water added in our watering regimes and the amount of water lost (demonstrated two times – Figure S7B, C - during watering regime of the grafted experiment, Figure S7A).

### Gas exchange measurements and water-use efficiency calculations

Gas exchange measurements including photosynthesis and transpiration rates, and stomatal conductance (via calculation), were obtained using a LI-COR 6400XT infrared gas analyzer (Lincoln, Nebraska, USA), both in the field and the glasshouse between 12:00 and 14:00 (‘PM’; Figure 6).

The LI-6400XT was combined with a Leaf Chamber Fluorometer in the glasshouse to obtain chlorophyll fluorescence measurements after 6 h of dark adaptation (lights off at 22:00, measurements from 4:00 to 6:00; ‘AM’; Figure 6B). A saturating pulse of light was applied to the dark-adapted leaves to ensure that all photosystem II (PII) energy was released as fluorescence and detected as the F_m_ value. F_v_ was calculated from F_m_ minus F_0_ (F_0_ being the base level of fluorescence emitted without the saturating pulse). Fv/Fm represents the maximum quantum yield of PII, which was used as a measure of stomatal-unrelated photosynthesis limitations (Signarbieux and Feller, 2011).

Water-use efficiency (WUE) was calculated as the ratio of photosynthetic rate (µmol CO_2_/m_2_s) to transpiration rate (mmol H_2_O/m_2_s), thus resulting in units of carbon dioxide molecules used per 1000 water molecules (Figure 6A, B, C).

### Micro-grafting

Seven-day-old seedlings were micro-grafted as described previously (Fragoso et al., 2011), with EV scions grafted to both EV (EV/EV) and irMPK4 (EV/irMPK4) rootstocks, and irMPK4 grafted only to irMPK4 (irMPK4/irMPK4) rootstocks. The average grafting success was 90% (p > 0.05 between genotypes, ANOVA, Tukey HSD *post hoc*).

### Transcript abundance

RNA was extracted with TRIzol reagent (Invitrogen) according to the manufacturer’s instructions. cDNA was synthesized from 500 ng of total RNA using RevertAid H Minus reverse transcriptase (Fermentas) and oligo (dT) primer (Fermentas). qPCR was performed in a Mx3005P PCR cycler (Stratagene) using SYBR GREEN1 kit (Eurogentec) using TaqMan primer pairs and double fluorescent dye-labeled probe. An *N. attenuata* sulfite reductase (*ECI*) was used as a standard housekeeping gene for normalization, and its primer sequences and probe, as well as the *MPK4* primer sequences and probes, are as published previously (Wu et al., 2007). Ct values were converted to relative transcript abundances using the standard curve method (Figure 2C, Figure S1). Silencing efficiency was calculated using the ΔΔCt method.

### Statistical analysis and pseudoreplication

All data were analyzed using R version 3.4.2 (RC Team 2017) and RStudio version 1.0.153 (RStudio Team 2016). Replication for experiments is indicated in the figure captions, including pseudoreplication. Pseudoreplication resulted from plants of the same genotype being measured from within the same population or pair: in the field experiment (Figure 3), the ‘top’ two plants of each mixed or monoculture MPxCC population were measured, while all four EVxCC plants in the 0% MPxCC populations were measured (Figure 3A; Figure S3B). In the glasshouse population experiment, all four central plants in all population types were measured (bottom, Figure 4A; Figure S5A). In the ungrafted paired experiment, both plants were measured in all pots, including the monocultures (Figure 5). In the grafted experiment, only the grafted plant was measured, so no pseudoreplication occurred.

Pseudoreplication was accounted for in our statistical analysis: these datasets were fit to linear mixed effect (LME) models with the population or pair they originated from indicated as a random effect. These models were checked for outliers, homoscedasticity and normality. Pairwise *post hoc* comparisons were made using the R package *emmeans* (Lenth et al., 2019).

Datasets without pseudoreplication were fit to the best suited of either AOV, LM, GLS or LME models, and were also checked for outliers, homoscedasticity and normality. Pairwise *post hoc* comparisons were extracted as above, or else using Tukey HSD tests following significant main effects in ANOVA.

## Acknowledgements

We thank the glasshouse team at the Max Planck Institute for Chemical Ecology and colleagues in the 2016 field work team for support; the technical staff at the Department of Molecular Ecology, particularly Dr. Klaus Gase, as well as Lucas Cortes Llorca, for advice on genetic material extractions; and the International Max Planck Research School (IMPRS) on the Exploration of Ecological Interactions with Chemical and Molecular Techniques and the Young Biodiversity Research Training Group - yDiv for their support of E.M. and H.V.

## Supplemental Figures and Tables

**Figure S1.**
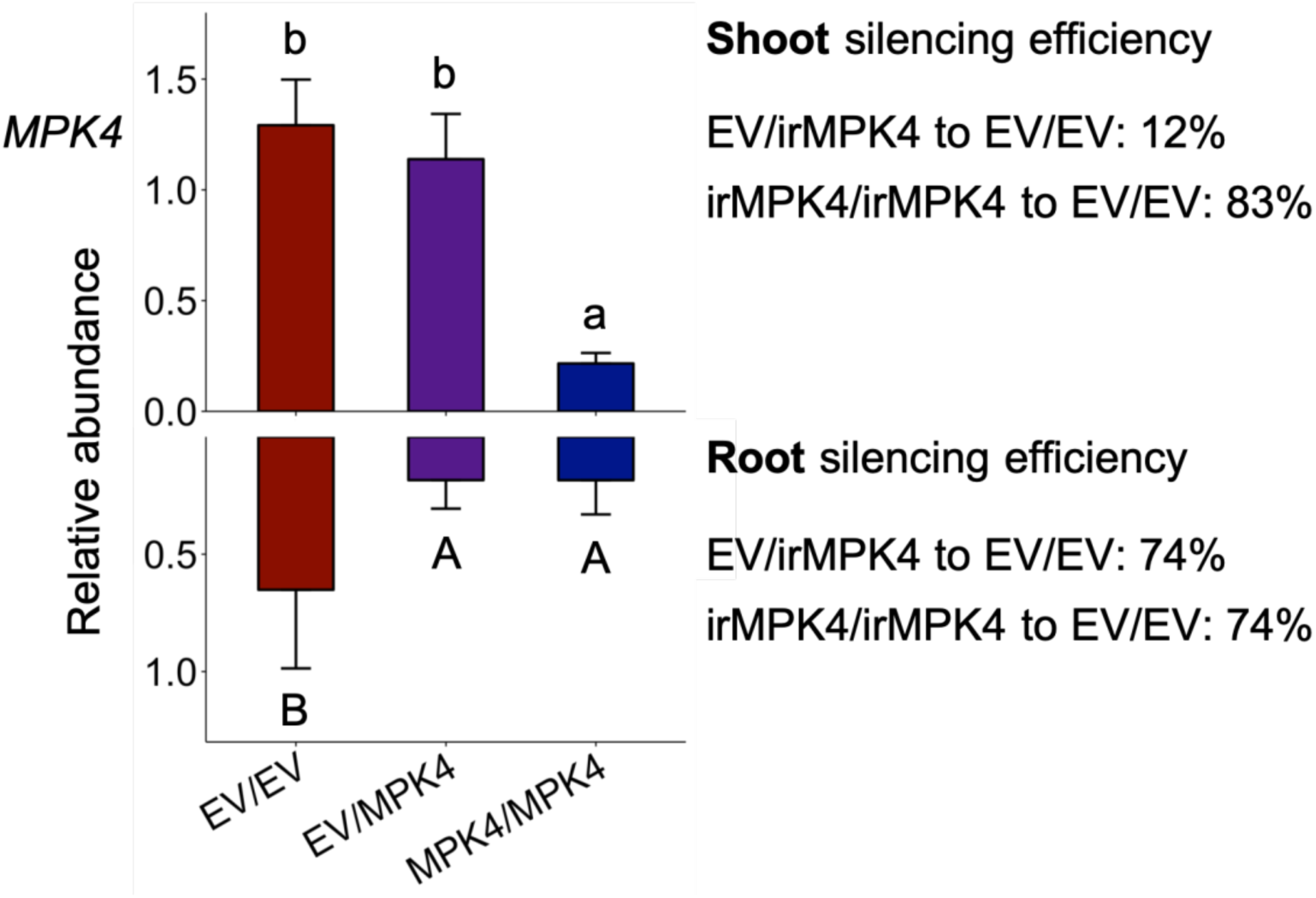
Relative transcript abundance (+ CI, n = 8 for EV/EV, 9 for EV/irMPK4, 6 for irMPK4/irMPK4) and silencing efficiency of *MPK4* in the shoots (top) and roots (bottom) of the glasshouse grafted pair experiment.

**Figure S2.**
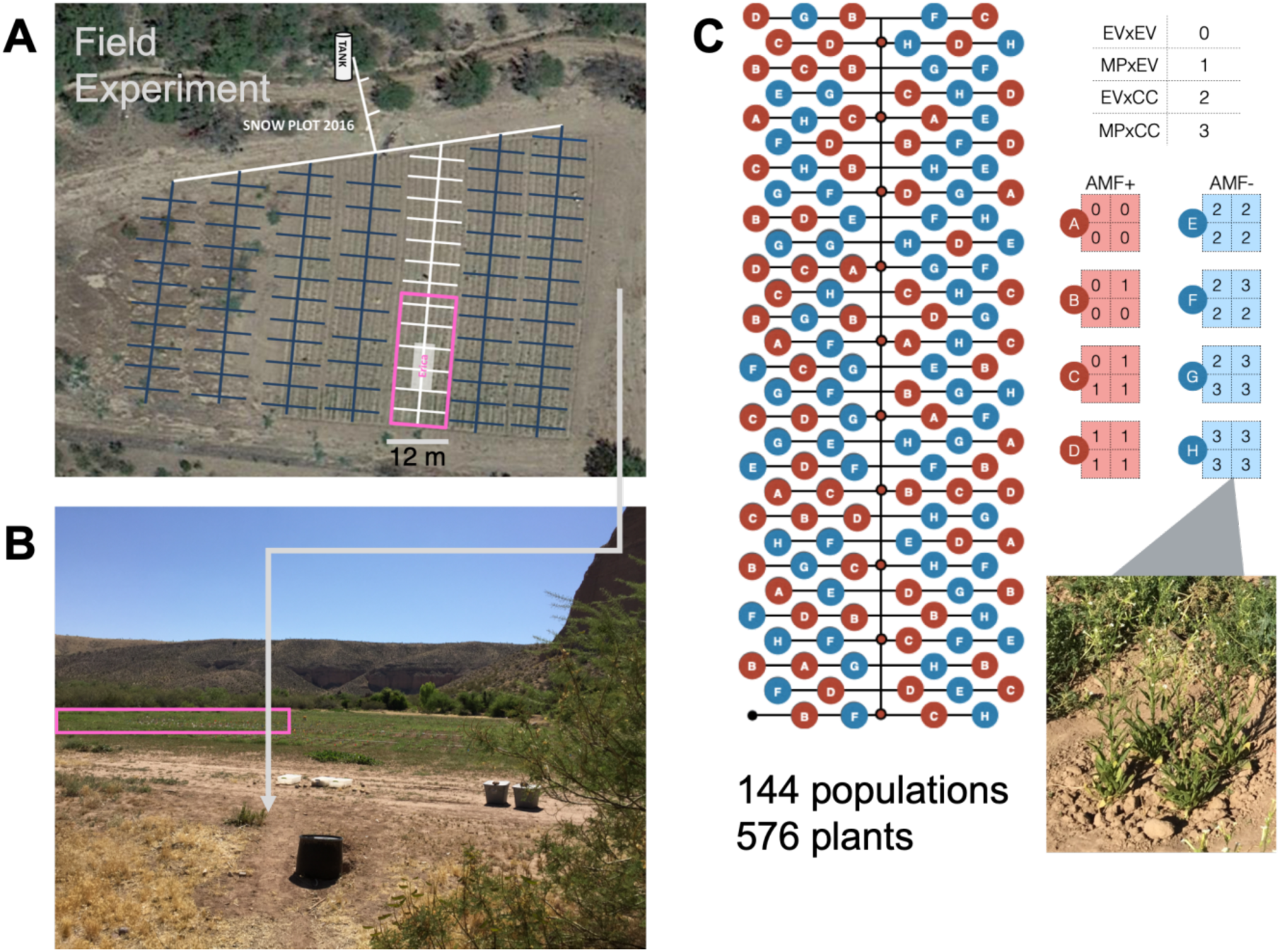
Layout of field population experiment at the Lytle Ranch Reserve field plot (“Snow Plot”, Utah) from an (A) aerial view (pink, Google Images), from a (B) side view (pink, picture by E.M.), and as a (C) schematic of all planted populations, with an example population at harvest, 53 days post planting (inset, picture by E.M).

**Figure S3.**
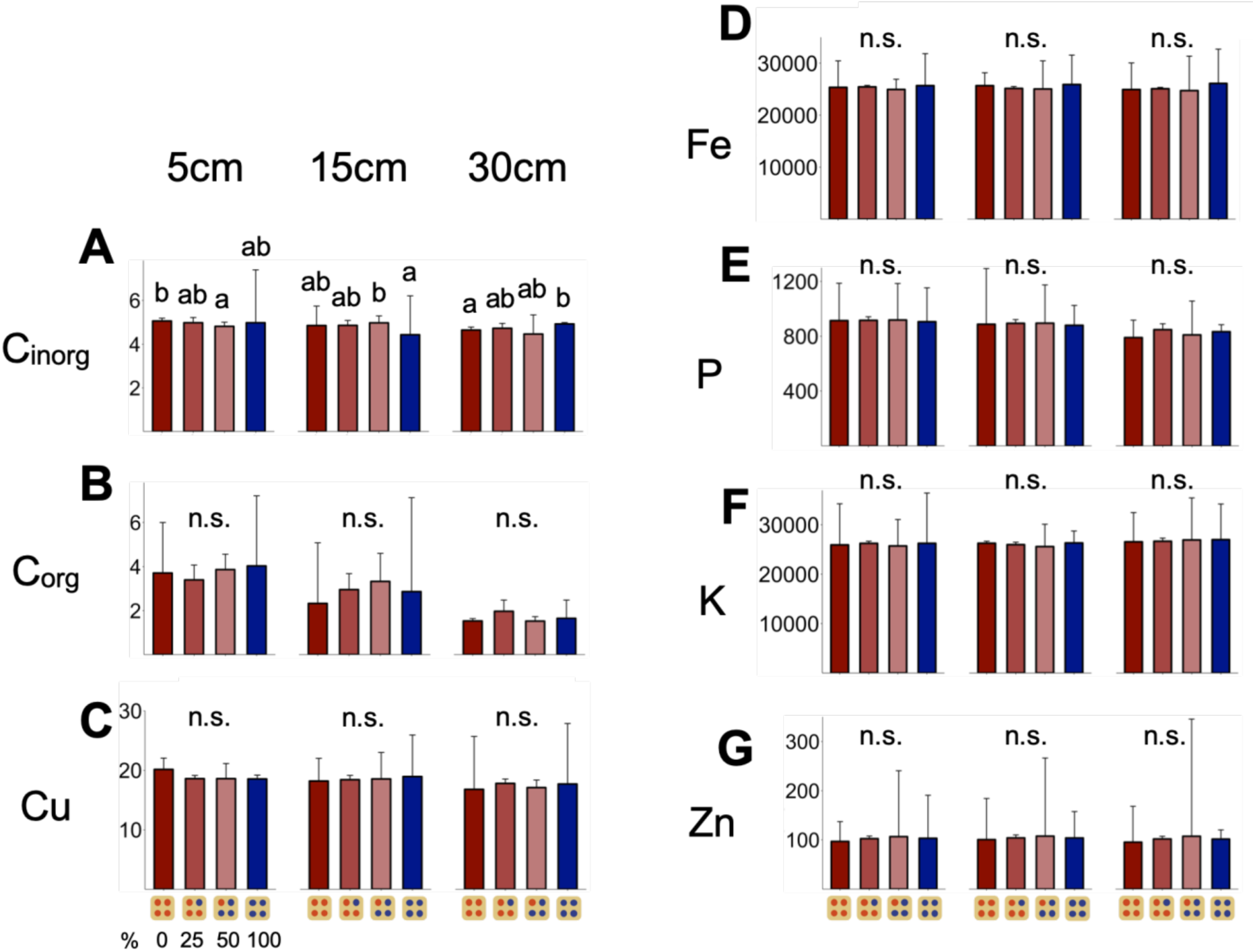
Element amounts (mg/kg + CI, y-axis, n = 3) of (A) inorganic carbon, (B) organic carbon, (C) copper, (D) iron, (E) phosphorus, (F) potassium, and (G) zinc, taken from soil cores 5, 15 and 30 cm below the center dripper of each population type (x-axis).

**Figure S4.**
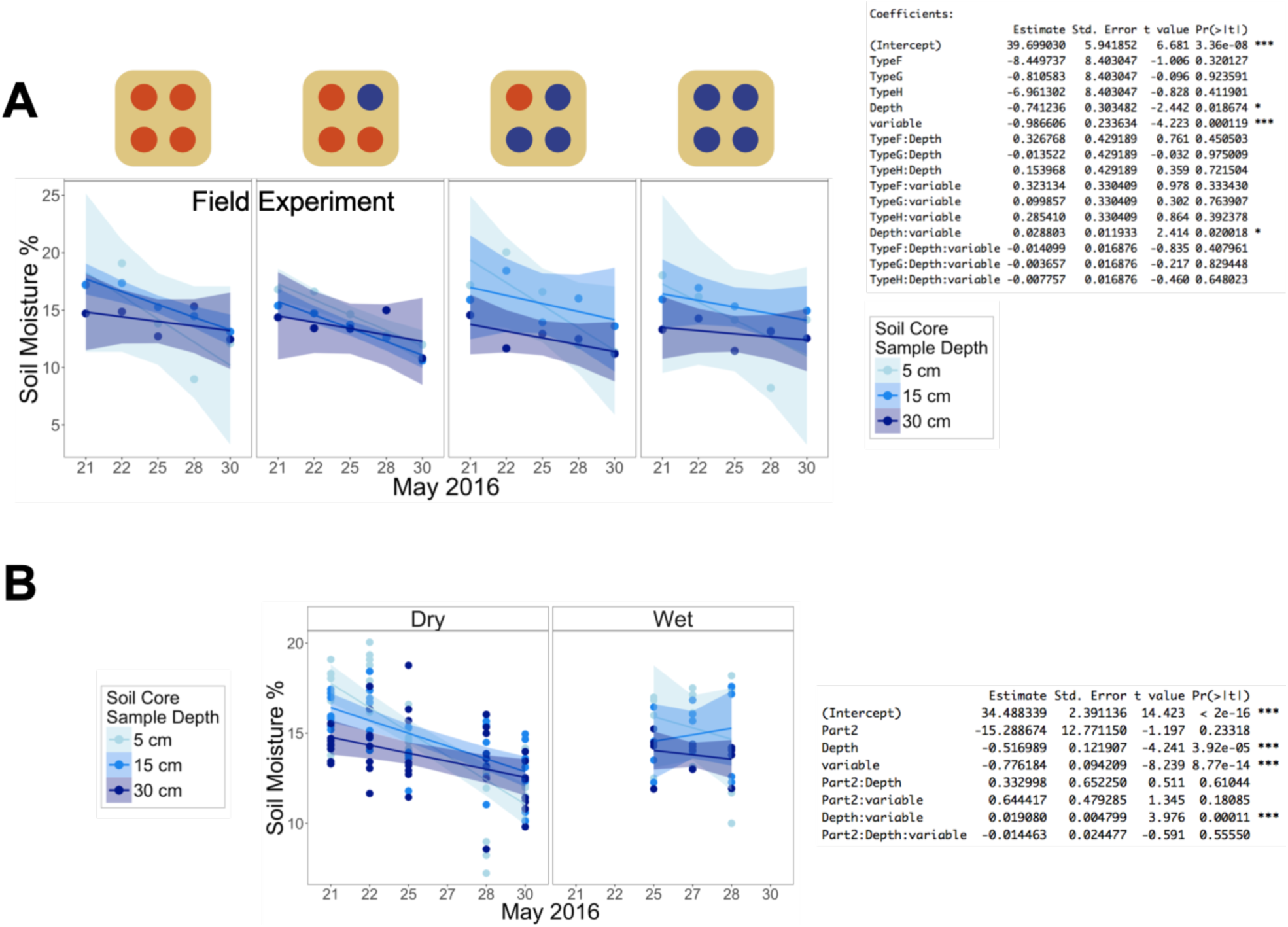
Soil moisture (%, y-axis, n = 1) of soil cores taken 5, 15 and 30 cm below the center dripper of each population type over 9 days (x-axis), either (A) cumulatively across the plot or (B) divided into the “Dry” and “Wet” subsections (see *Water treatments* in Materials and Methods), with corresponding regression analysis results.

**Figure S5.**
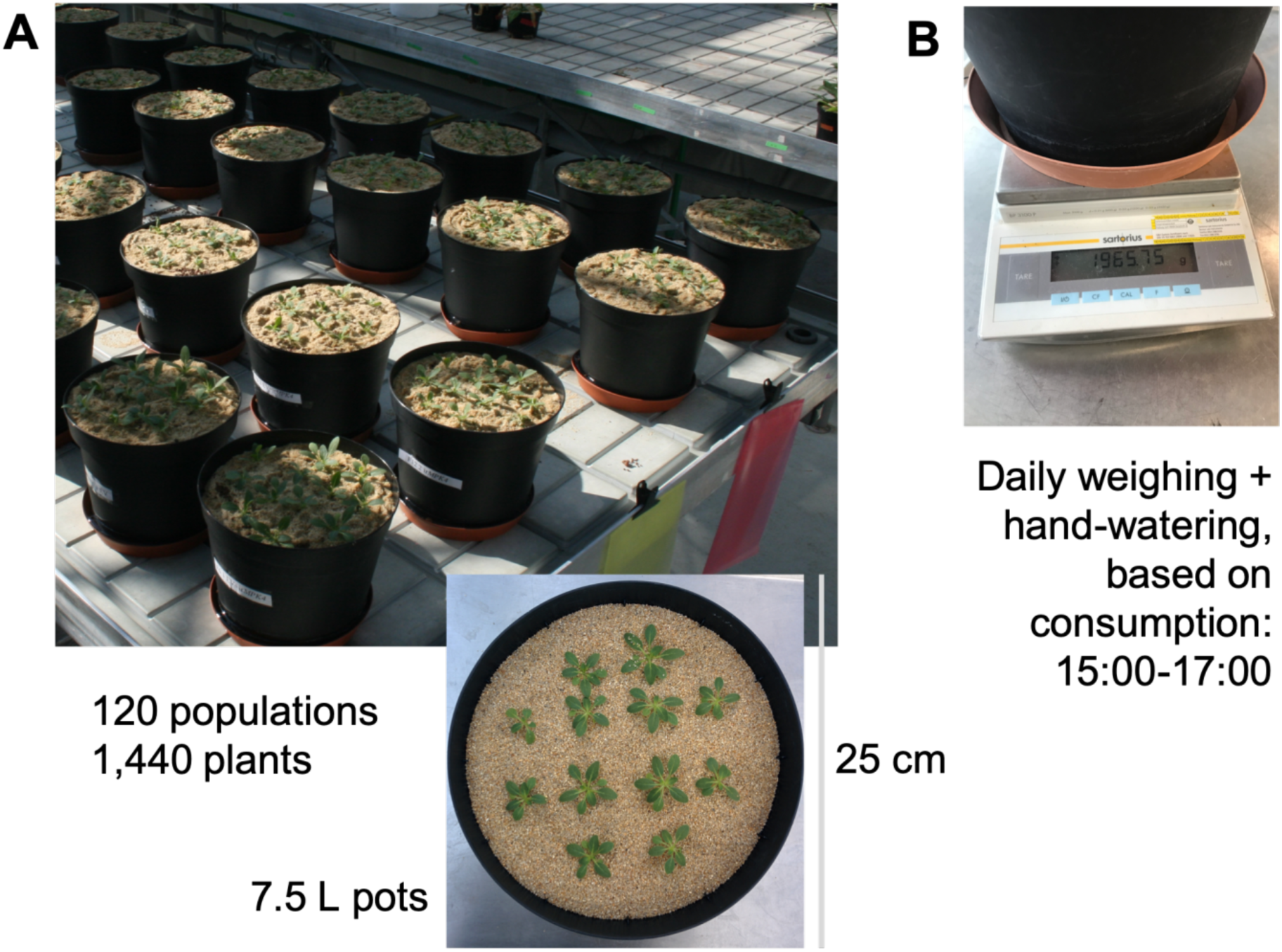
Layout of glasshouse population experiment as pictured from (A) above the glasshouse table (picture by D.M), and above a single example pot (inset, picture by E.M.); each 7.5L pot was (B) weighed daily to determine how much water was lost from the day before as well as provide consumption based watering based on daily water loss (picture by E.M.).

**Figure S6.**
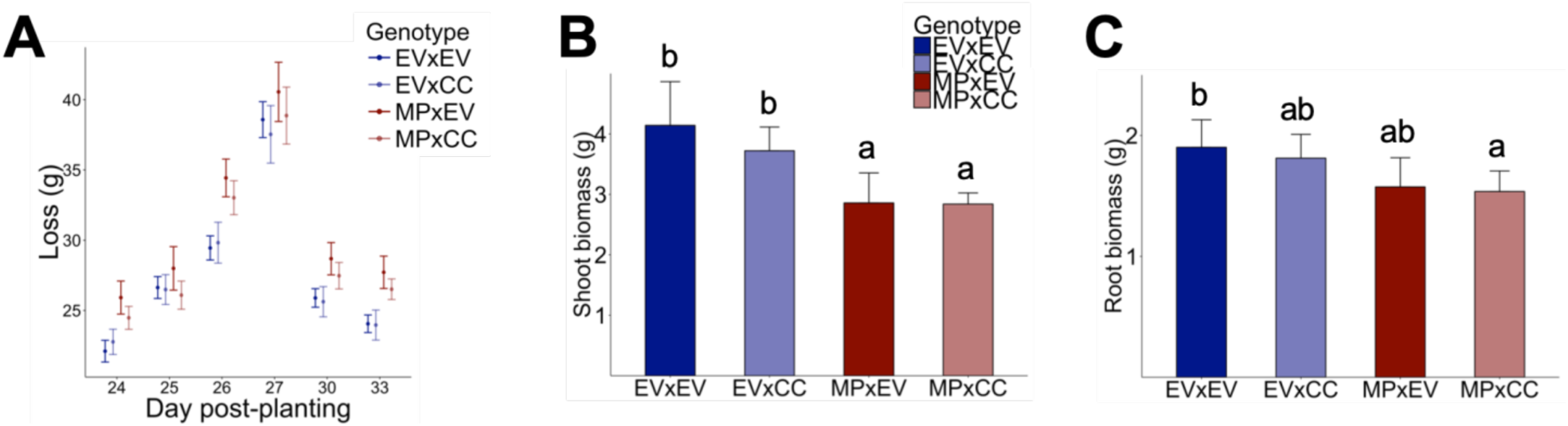
Cross characterization experiment comparing EVxEV to EVxCC and MPxEV to MPxCC in (A) water loss per genotype (g ± CI, n = 15-16, y-axis) per day (x-axis), (B) fresh shoot biomass (g + CI, n = 5-7) per genotype (x-axis), and (c) fresh root biomass (g + CI, n = 6-7).

**Figure S7.**
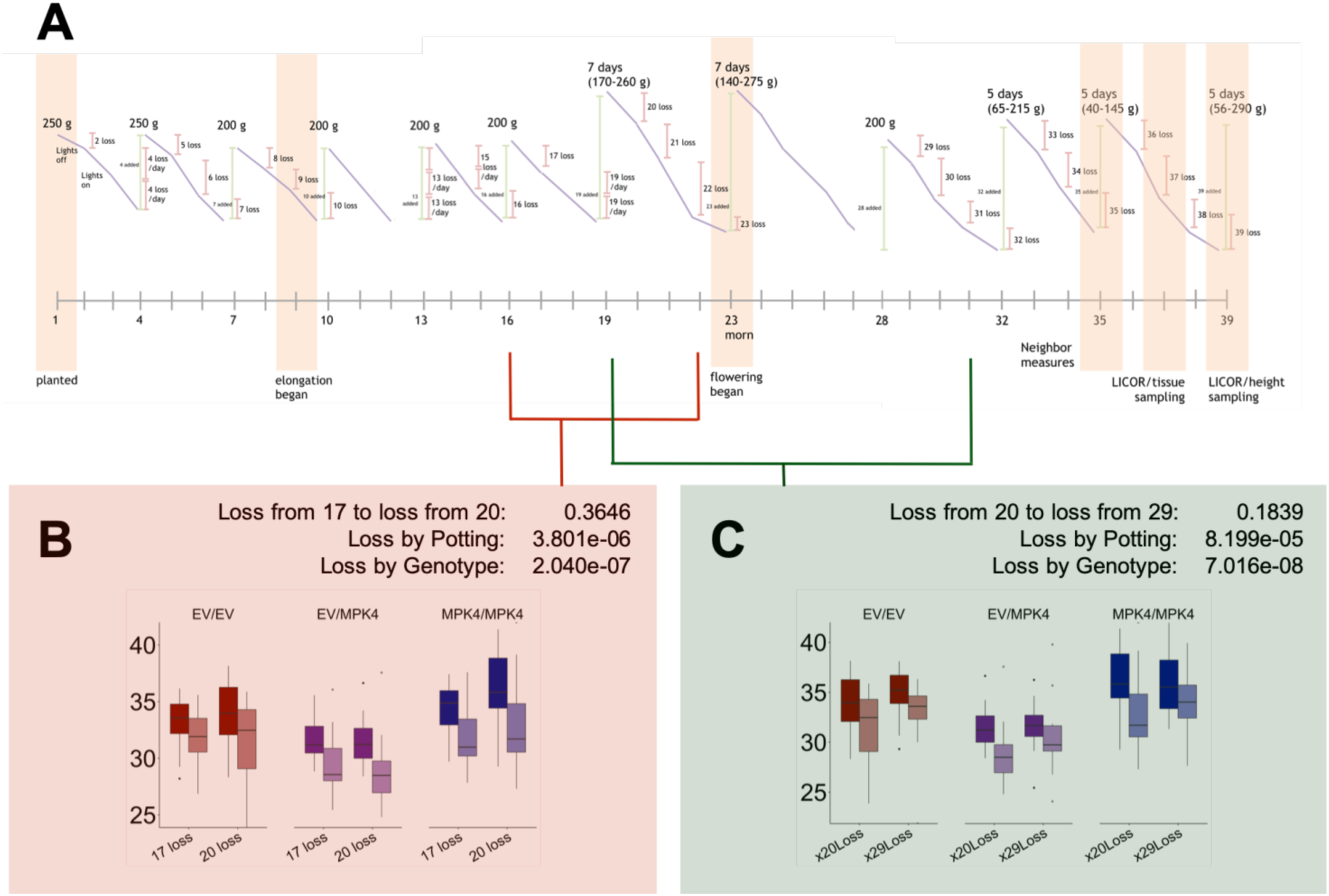
Watering regime of the glasshouse grafted pair experiment (A), with peak text indicating water (g) provided to pots on the respective day of the experiment (x-axis) and orange highlights indicating key experimental events (developmental changes and sampling times). Two analyses on whether (B) increasing water given to a pot changes the water loss per day of the pot (y-axis) from experimental day 17 (given 200 g water) to 20 (given 170-260 g water, x-axis) or whether (C) decreasing water given to a pot changes the water loss per day of the pot (y-axis) from experimental day 20 (given 170-260 g water) to 29 (given 200 g water, x-axis), are each accompanied with respective p-values from an ANOVA conducted on the variables: loss between days, loss between potting type (single: dark colors; paired: faded colors), or loss by genotype of the grafted plant in the pot (red: EV/EV; purple: EV/irMPK4; blue: irMPK4/irMPK4).

**Table S1.** Statistical *emmeans* contrasts of hemizygous to homozygous crosses for Figure S6A.

| 1108 | DPP | Contrast | t value | p value |
| --- | --- | --- | --- | --- |
| 1109 |  |  |  |  |
| 1110 | 24 | EVxEV<sub>[N=16]</sub> to EVxCC<sub>[N=16]</sub> | 0.635 | 0.9208 |
| 1111 |  | MPxEV<sub>[N=15]</sub> to MPxCC<sub>[N=16]</sub> | 0.673 | 0.9072 |
| 1112 | 25 | EVxEV<sub>[N=16]</sub> to EVxCC<sub>[N=16]</sub> | -1.799 | 0.2757 |
| 1113 |  | MPxEV<sub>[N=15]</sub> to MPxCC<sub>[N=16]</sub> | 1.034 | 0.7298 |
| 1114 | 26 | EVxEV<sub>[N=16]</sub> to EVxCC<sub>[N=16]</sub> | 1.176 | 0.6427 |
| 1115 |  | MPxEV<sub>[N=15]</sub> to MPxCC<sub>[N=16]</sub> | 0.640 | 0.9189 |
| 1116 | 27 | EVxEV<sub>[N=16]</sub> to EVxCC<sub>[N=16]</sub> | -2.342 | 0.0906 |
| 1117 |  | MPxEV<sub>[N=15]</sub> to MPxCC<sub>[N=15]</sub> | 0.063 | 0.9999 |
| 1118 | 30 | EVxEV<sub>[N=16]</sub> to EVxCC<sub>[N=15]</sub> | -0.472 | 0.9652 |
| 1119 |  | MPxEV<sub>[N=15]</sub> to MPxCC<sub>[N=16]</sub> | 0.690 | 0.9007 |
| 1120 | 33 | EVxEV<sub>[N=16]</sub> to EVxCC<sub>[N=15]</sub> | -0.389 | 0.9800 |
| 1121 |  | MPxEV<sub>[N=15]</sub> to MPxCC<sub>[N=16]</sub> | 0.105 | 0.9996 |
| 1122 |  |  |  |  |

DPP: days post-planting

